# A mitotic restriction point for extrachromosomal DNA hubs in cancer cells

**DOI:** 10.64898/2026.10.08.757707

**Authors:** Xiaowei Yan, Ivy Tsz-Lo Wong, Muhammad Murtaza Hassan, Rui Li, Katerina Kraft, Aditi Gnanasekar, Paul S. Mischel, Howard Y. Chang

## Abstract

Extrachromosomal DNA (ecDNA) is widespread in human cancers and promotes tumour evolution, intratumour heterogeneity and poor clinical outcomes^1–5^. ecDNA molecules tend to cluster into micron-scale nuclear hubs that enhance oncogene transcription^6^. However, how these higher-order assemblies form and persist remains largely unknown. Here we discover mitosis as the critical window for ecDNA hub organisation across cell generations. We develop an optical screen that identifies the mitotic-entry regulator WEE1 as an important regulator of hub integrity and oncogene expression. Analysis of WEE1 perturbation reveals that CDK1- and CDK2-regulated programmes cooperate to carry unresolved damage into mitosis, promoting reorganisation of ecDNA tethered to mitotic chromosomes. Long-term live-cell imaging shows that, in unperturbed cells, ecDNA hub architecture is remodelled primarily during mitosis and subsequently maintained throughout interphase. This coupling between mitotic and interphase organisation is conserved across diverse cancer types and ecDNA-associated oncogenes. Together, these findings define a mitotic restriction point that governs ecDNA hub organisation, oncogene expression and ecDNA inheritance through successive cell generations, revealing a potential window for therapeutic intervention in ecDNA-driven cancers.

---

Extrachromosomal DNA (ecDNA) has emerged as a prevalent form of oncogene amplification in human cancers, creating a dynamic genome architecture that can accelerate tumour evolution^1–5^. These megabase-sized circular DNA elements reside outside canonical chromosomes and function as autonomous oncogene carriers, often accumulating to hundreds of copies per cancer cell and driving high oncogene expression^7,8^. We previously showed that ecDNA molecules do not remain spatially isolated, but instead cluster into nuclear hubs that enhance transcriptional activity through intermolecular enhancer sharing^6^ and promote the coordinated inheritance of distinct ecDNA species^9^. Together, these findings suggest that ecDNA is governed by higher-order spatial regulation within the nucleus. One possibility is that ecDNA hubs are dynamic assemblies that continuously form and dissolve throughout the cell cycle. Alternatively, hub organisation may be controlled at discrete restriction points, analogous to the G1 restriction point of the cell division cycle. To date, the mechanisms that maintain ecDNA hubs and the broader principles governing ecDNA spatial organisation remain poorly understood.

Progress towards resolving these questions has been limited by the lack of approaches to systematically perturb ecDNA spatial organisation, particularly hub formation. This challenge is compounded by the intrinsic heterogeneity of ecDNA, which makes conventional screening difficult. To overcome this barrier, we developed an optical screening approach that enabled systematic interrogation of ecDNA organisation and identification of perturbations that disrupt hub integrity. Combining this platform with long-term live-cell imaging, we uncovered unexpected regulators of ecDNA hubs and defined a temporal framework for their spatial control. These analyses revealed that ecDNA hub organisation is not continuously re-established during interphase, but is instead strongly constrained by events in the preceding mitosis. Our findings identify mitosis as a key restriction point for ecDNA hub reorganisation and reveal a spatial inheritance rule that governs ecDNA organisation across cell generations.

## hiFISH-96 screening identifies WEE1

To systematically identify perturbations of ecDNA hub organisation, we established hiFISH-96, a 96-well high-content DNA FISH platform coupled to automated single-cell image analysis (Fig. 1a; Extended Data Fig. 1). We applied this strategy to COLO320DM cells, an ecDNA-positive colorectal cancer model harbouring *MYC*-amplified ecDNAs that frequently assemble into prominent nuclear hubs^6^. We quantified ecDNA organisation on an object-by-object basis and extracted spatial features, including ecDNA dispersal, measured as spatially resolved foci per cell, and radial distribution.

**Fig. 1.**
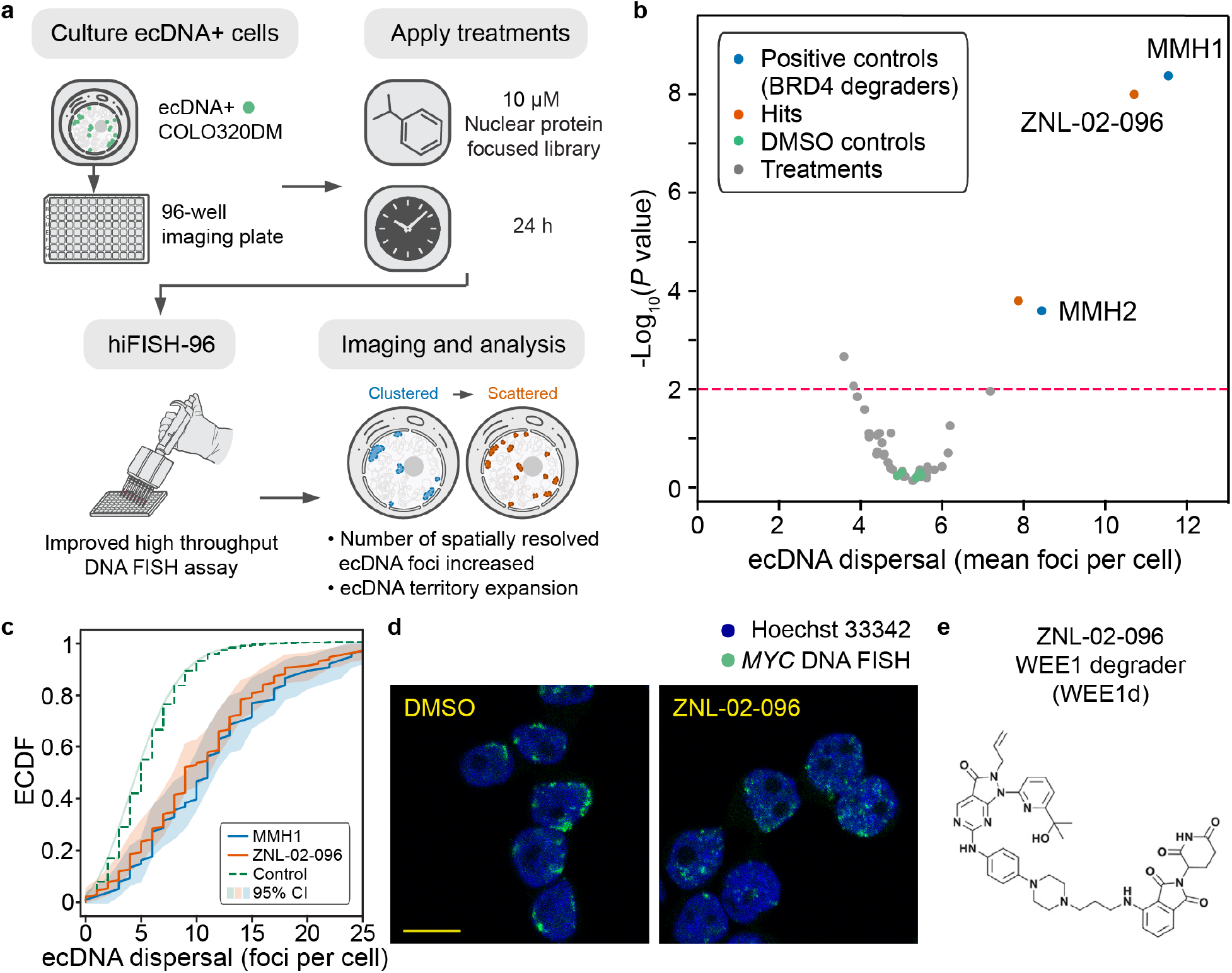
hiFISH-96 identifies a WEE1 degrader that disperses ecDNA hubs. **a,** Schematic of the hiFISH-96 workflow for high-content DNA FISH screening and automated single-cell quantification of ecDNA spatial organisation in COLO320DM cells. A candidate-based library of 43 small molecules targeting nuclear proteins was screened at 10 μM. **b,** Screening plot showing ecDNA dispersal, quantified as the mean number of spatially resolved ecDNA foci per cell, and the associated *P* value for each of the 39 evaluable compounds and DMSO controls. Four compounds causing substantial cell death were excluded. The BRD4 degraders MMH1 and MMH2, included as positive controls^31,32^, are shown in blue; screen hits are orange, DMSO controls are green and the remaining treatments are grey. The dashed line denotes *P* = 0.01. **c,** Empirical cumulative distribution functions (ECDFs) of ecDNA dispersal, measured as foci per cell, in the hiFISH-96 screen following treatment with ZNL-02-096 (orange), MMH1 (blue) or DMSO (green). The DMSO distribution comprises cells pooled across all DMSO control wells. Shading denotes 95% confidence intervals. **d,** Representative single-z-plane DNA FISH images from the hiFISH-96 screen showing *MYC*-containing ecDNA (green) and Hoechst 33342-stained nuclei (blue) in DMSO- and ZNL-02-096-treated COLO320DM cells. Scale bar, 10 μm. **e,** Chemical structure of the WEE1 degrader ZNL-02-096 (WEE1d). Statistical significance in **b** was assessed using two-sample Kolmogorov-Smirnov tests with repeated subsampling. For each compound, up to 100 nuclei were randomly sampled and compared with an equal number sampled from the pooled DMSO controls. Sampling was repeated 50 times, and the plotted value represents the mean −log10(*P*) across iterations. This approach made statistical significance comparable across conditions with different numbers of segmented nuclei.

To define robust screening metrics, we performed repeated DMSO control experiments and assessed the reproducibility of ecDNA organisational features across independent hiFISH-96 runs. Several metrics showed stable behaviour under baseline conditions, including ecDNA dispersal, quantified as the number of spatially resolved foci per cell, which was selected as the primary screening metric (Extended Data Fig. 2). Because ecDNA hubs are nuclear assemblies, we reasoned that their organisation might depend on proteins regulating chromatin and other nuclear processes. We therefore applied hiFISH-96 in a candidate-based screen of 43 small molecules targeting nuclear proteins. Four compounds induced substantial cell death and were not further evaluated. The remaining 39 compounds were evaluated for their ability to increase ecDNA dispersal relative to DMSO-treated cells (Fig. 1b,c; Supplementary Data 1). As expected, two BRD4 degraders included as positive controls scored strongly, consistent with previous work identifying BRD4 as a regulator of ecDNA hub integrity in COLO320DM^6,10^. Among the screened compounds, ZNL-02-096 emerged as the strongest hit, producing a marked increase in ecDNA dispersal (Fig. 1b,c).

Representative images from the screen showed that ZNL-02-096 disrupted compact ecDNA hubs and produced a more dispersed ecDNA pattern (Fig. 1d). In addition to increasing ecDNA dispersal, ZNL-02-096 and BRD4 degraders also shifted ecDNA radial distribution inward, indicating that hub dispersal is accompanied by broader reorganisation of ecDNA spatial positioning (Extended Data Fig. 3). ZNL-02-096 is a recently developed WEE1 degrader^11^, which we refer to hereafter as WEE1d (Fig. 1e). Thus, hiFISH-96 identified the WEE1 degrader ZNL-02-096 as a compound that strongly perturbs ecDNA hub organisation.

## WEE1 degradation disperses ecDNA hubs

We therefore examined whether WEE1 loss was responsible for the ecDNA dispersal phenotype identified in the screen. WEE1d induced dose-dependent ecDNA dispersal, and immunoblotting confirmed robust WEE1 depletion at concentrations that elicited this phenotype; based on these titrations, 2 μM WEE1d was used for subsequent analyses (Extended Data Fig. 4a,b). We next compared WEE1d with MK-1775, a well-characterised and widely used WEE1 inhibitor (WEE1i)^12^. As predicted, pharmacological WEE1 inhibition phenocopied WEE1d-induced ecDNA dispersal (Fig. 2a,b) and produced a similar dose-dependent response (Extended Data Fig. 4c), supporting an on-target effect of WEE1d. Because targeted protein degraders act by recruiting their target proteins to the ubiquitin-proteasome system, we then asked whether WEE1d-induced dispersal depended on proteasome activity. Consistent with this mechanism, proteasome inhibition with MG132^13^ blocked the dispersal phenotype (Fig. 2c,d), further supporting a degradation-dependent effect of WEE1 loss.

**Fig. 2.**
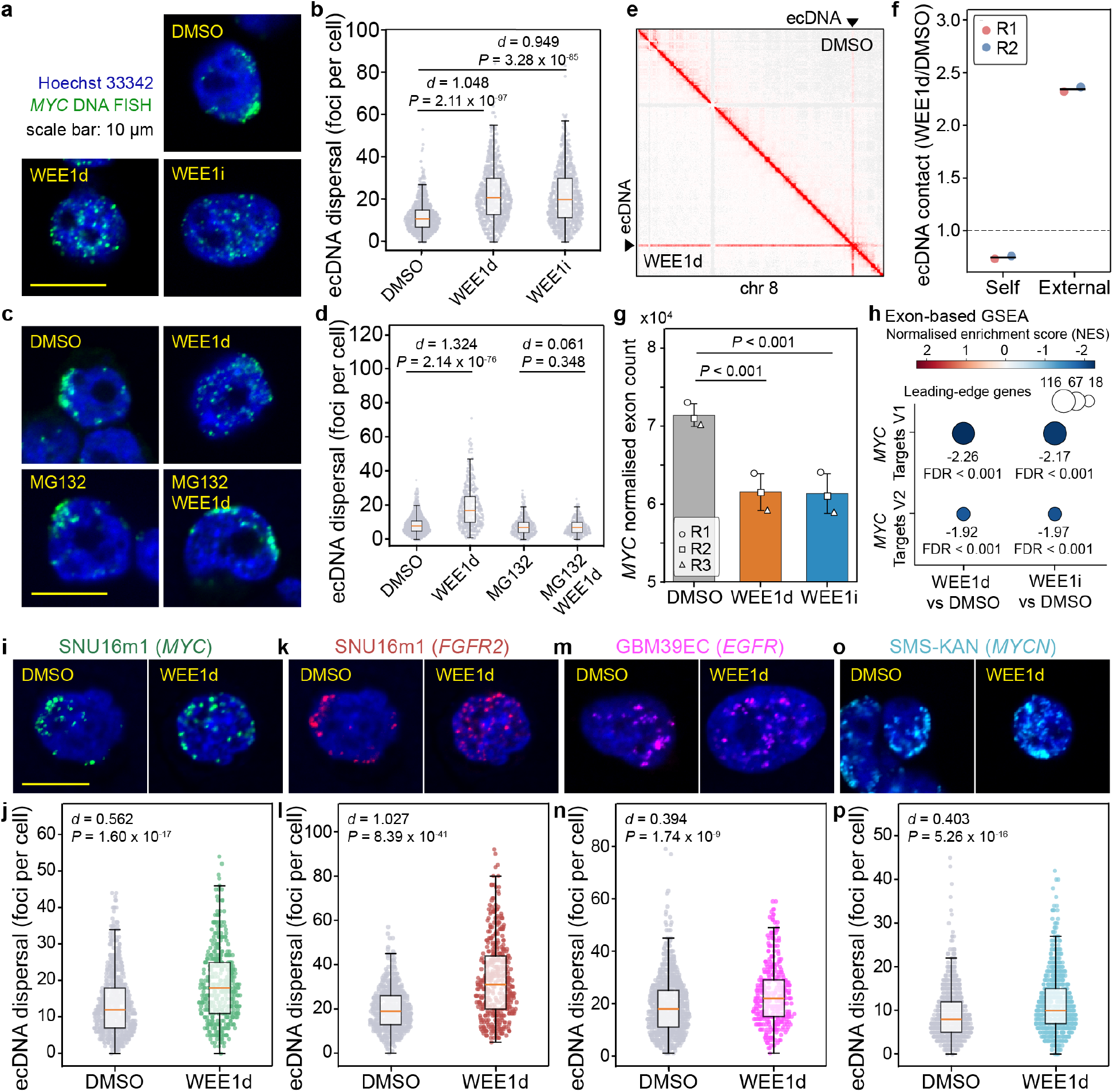
WEE1 perturbation disperses ecDNA hubs and reduces oncogene expression. **a,b,** Representative DNA FISH images (**a**) and ecDNA dispersal, quantified as foci per cell (**b**), in COLO320DM cells treated for 24 h with DMSO, WEE1d or WEE1i (MK-1775) (*n* = 4 biological replicates). **c,d,** Representative DNA FISH images (**c**) and ecDNA dispersal (**d**) after 24 h treatment with DMSO, WEE1d, MG132 or WEE1d plus MG132 (*n* = 3 biological replicates). **e,f,** Chromosome 8 Hi-C maps from DMSO- and WEE1d-treated COLO320DM cells (**e**; DMSO, upper triangle; WEE1d, lower triangle) and WEE1d-to-DMSO ratios of ecDNA self-contacts and contacts with external chromosomal regions (**f**; *n* = 2 biological replicates). Symbols denote biological replicates and horizontal lines indicate means. **g,h,** RNA-seq analyses after 24 h treatment with DMSO, WEE1d or WEE1i (*n* = 3 biological replicates): DESeq2-normalised *MYC* exon counts (**g**; mean ± s.d.; symbols denote biological replicates) and exon-based GSEA of Hallmark *MYC* Targets V1 and V2 (**h**). Colour indicates normalised enrichment score (NES), and bubble size indicates the number of leading-edge genes. NES and FDR *q* values are shown. **i–p,** Representative DNA FISH images and ecDNA dispersal for *MYC*- and *FGFR2*-containing ecDNA in SNU16m1 cells treated for 24 h (**i–l**), *EGFR*-containing ecDNA in GBM39EC cells treated for 72 h (**m,n**) and *MYCN*-containing ecDNA in SMS-KAN cells treated for 24 h (**o,p**). Representative DNA FISH images show single z-planes. FISH signals are coloured as indicated; Hoechst 33342 is blue. Concentrations: 2 μM WEE1d, 500 nM WEE1i and 10 μM MG132. Sample sizes are provided in Supplementary Table 1. Two-sided Welch’s *t*-tests were used in **b,d,j,l,n,p**; absolute Cohen’s *d* and exact *P* values are shown. DESeq2 Wald tests with Benjamini-Hochberg adjustment were used in **g**; adjusted *P* values are shown. Box plots show medians and interquartile ranges; whiskers extend to 1.5× the interquartile range. Scale bars, 10 μm.

To further assess the impact of WEE1 loss on ecDNA spatial organisation, we examined chromatin contacts genome-wide by Hi-C. Comparison of WEE1d-treated and DMSO-treated COLO320DM cells revealed reduced self-interaction among ecDNAs and a reciprocal increase in contacts between ecDNA and external chromosomal loci, a redistribution confirmed by quantitative analysis (Fig. 2e,f). These orthogonal measurements support the conclusion that WEE1 loss disperses ecDNA hubs and broadens the chromatin-contact landscape of ecDNA.

Previous work showed that ecDNA hubs enhance oncogene transcription through intermolecular interactions that facilitate the sharing of regulatory elements, including enhancers, between ecDNA molecules^6^. To determine whether WEE1d-induced hub dispersal was accompanied by functional changes, we performed RNA sequencing following WEE1 degradation or inhibition. Both perturbations markedly reduced the steady-state expression of ecDNA-encoded *MYC* (Fig. 2g), and gene-set enrichment analysis revealed coordinated downregulation of *MYC* target programmes (Fig. 2h; Extended Data Fig. 5a,b). Analysis of intronic reads, used as a proxy for nascent transcription, showed a concordant trend towards reduced *MYC* transcription and negative enrichment of *MYC* target programmes (Extended Data Fig. 5c–f). Thus, the structural disruption of ecDNA hubs following WEE1 perturbation coincides with reduced expression of ecDNA-encoded *MYC* and attenuation of its downstream transcriptional programmes.

We next asked whether this phenotype was restricted to COLO320DM cells or to *MYC*-amplified ecDNA. Across additional ecDNA-positive models, including SNU16m1 gastric cancer cells (*MYC* and *FGFR2* amplified on distinct ecDNAs), GBM39EC glioblastoma neurospheres (*EGFR* amplified on ecDNA) and SMS-KAN neuroblastoma cells (*MYCN* amplified on ecDNA), WEE1 degradation likewise increased ecDNA dispersal and reduced hub compaction (Fig. 2i–p). The magnitude of dispersal varied among models, potentially reflecting differences in basal ecDNA clustering and nuclear dimensions. Nevertheless, the response was observed across different cancer types and ecDNAs carrying distinct oncogenes, including *MYC*, *FGFR2*, *EGFR* and *MYCN*, indicating that WEE1-dependent control of ecDNA hub organisation is not restricted to a particular cancer type or ecDNA species. Together, these findings identify WEE1 as a shared regulator of ecDNA spatial organisation across diverse human cancer models.

## Mitosis drives hub reorganisation

We focused on COLO320DM cells for subsequent mechanistic studies because this model forms prominent ecDNA hubs that can be readily visualised and quantified. Beyond ecDNA hub dispersal, a prominent response to WEE1 degradation was altered mitotic progression. Given the established role of WEE1 in regulating the G2/M transition^14^, we characterised cell-cycle dynamics following WEE1 degradation in this ecDNA-positive model (Fig. 3a). Live-cell imaging using a FUCCI(CA) reporter^15^ showed that WEE1 degradation shortened the interval between G1 exit and mitotic entry and prolonged M phase approximately eightfold (Extended Data Fig. 6a–c), consistent with premature mitotic entry followed by delayed mitotic progression. To determine the fate of WEE1d-treated cells, we tracked 50 cells that were in interphase at the beginning of a 24-h imaging period (Fig. 3b). Of these, 39 cells (78%) entered mitosis. Among them, 28 (71.8%) completed mitosis and generated progeny: 26 (66.7%) remained alive and trackable until the end of imaging, 1 (2.6%) died 4 h after returning to interphase and 1 (2.6%) was lost to tracking 8.5 h after returning to interphase. Of the remaining cells, 10 (25.6%) remained arrested in mitosis at the end of imaging and only 1 (2.6%) died during mitosis (Fig. 3b). Thus, although WEE1 inhibition is frequently associated with mitotic catastrophe^16^, mitotic death was rare in WEE1d-treated COLO320DM cells under these conditions. Among the 40 trackable cells in interphase at 24 h, 30 (75%) had passed through mitosis during imaging (Fig. 3b), indicating that the endpoint interphase population consisted predominantly of cells that had experienced an aberrant mitosis. These observations raised the possibility that WEE1d-induced ecDNA hub dispersal arises during aberrant mitosis and subsequently becomes apparent in interphase.

**Fig. 3.**
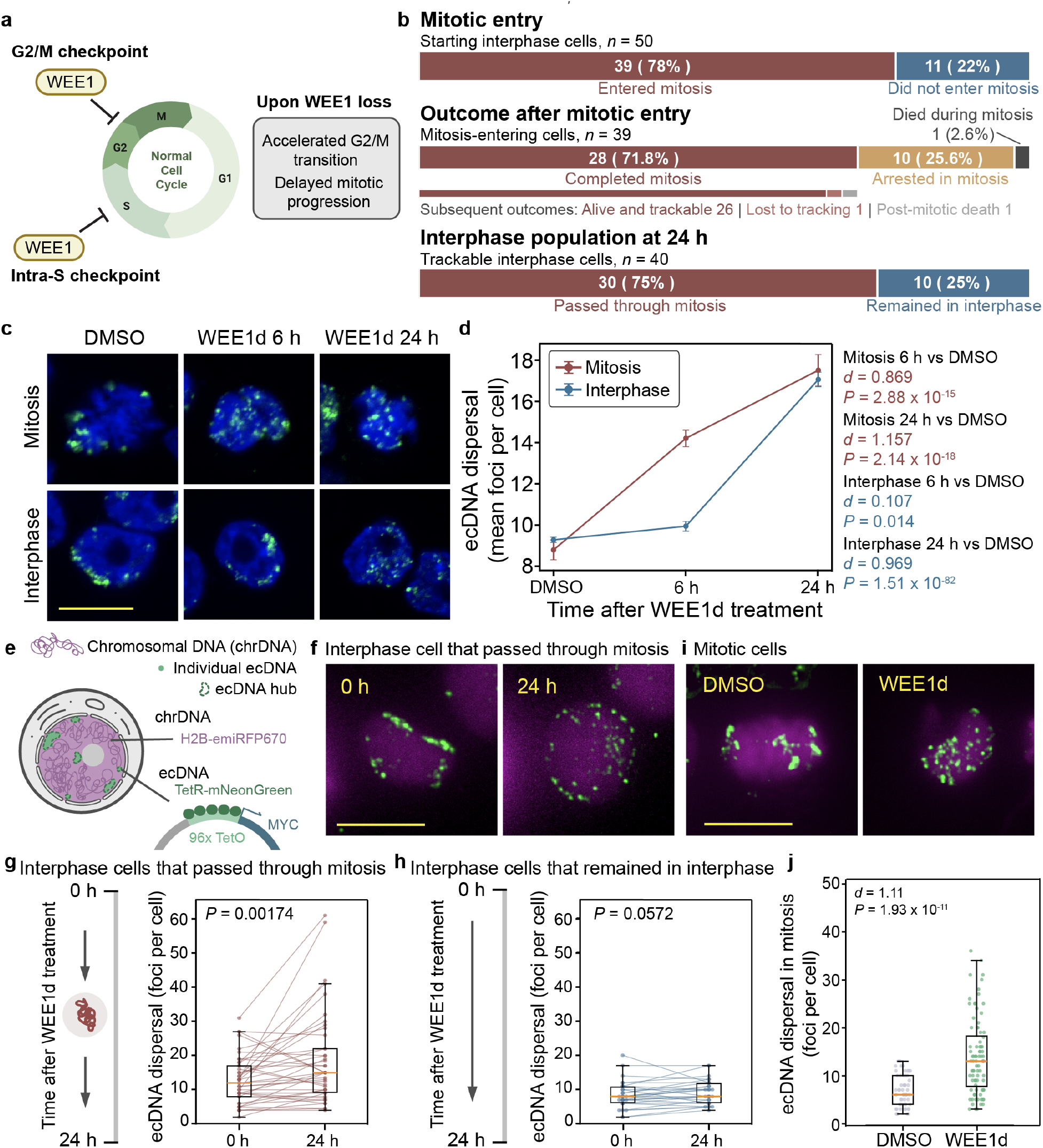
WEE1 degradation accelerates mitotic entry and drives mitosis-dependent ecDNA hub dispersal. **a,** Schematic of WEE1 functions in the intra-S and G2/M checkpoints and the effects of WEE1 loss. **b,** Mitotic entry and cell outcomes during 24 h of FUCCI(CA) tracking following WEE1 degradation (*n* = 2 biological replicates). Cells in interphase at movie onset were tracked to determine the fraction entering mitosis and the outcomes of mitosis-entering cells. The endpoint analysis shows the mitotic history of cells in interphase at 24 h. **c,d,** Representative DNA FISH images (**c**) and ecDNA dispersal, quantified as mean foci per cell (**d**), in interphase and mitotic COLO320DM cells treated with DMSO or WEE1d for 6 or 24 h (*n* = 3 biological replicates). **e,** Live-cell ecDNA-labelling strategy: a 96× TetO array adjacent to ecDNA-encoded *MYC* was visualised with TetR-mNeonGreen, and chromosomal DNA with H2B-emiRFP670. **f–h,** Representative images of a WEE1d-treated interphase cell that passed through mitosis (**f**) and paired measurements of ecDNA dispersal in interphase cells at 0 and 24 h, grouped according to whether they passed through mitosis (**g**) or remained in interphase throughout imaging (**h**), pooled across *n* = 4 biological replicates. **i,j,** Representative mitotic-cell images (**i**) and ecDNA dispersal (**j**) under DMSO or WEE1d treatment, pooled across *n* = 4 biological replicates. In **c**, *MYC* DNA FISH is green and Hoechst 33342 is blue; in **f,i**, ecDNA is green and chromosomal DNA is magenta. **c** shows single-z-plane images; **f,i** show SVCC-rendered maximum-intensity projections. WEE1d was used at 2 μM. Sample sizes are provided in the figure and Supplementary Table 1. Two-sided Welch’s *t*-tests were used in **d,j**, with absolute Cohen’s *d* reported. In **d**, points show means and error bars denote s.e.m. Paired two-sided Wilcoxon signed-rank tests were used in **g,h**. Exact *P* values and effect sizes are shown. Box plots show medians and interquartile ranges; whiskers extend to 1.5× the interquartile range. Scale bars, 10 μm.

To resolve this temporal order, we quantified ecDNA organisation separately in mitotic and interphase cells using fixed-cell DNA FISH. By 24 h, when lineage tracking indicated that the interphase population consisted predominantly of cells that had passed through mitosis, ecDNA dispersal was evident in both populations (Fig. 3c,d). At the earlier 6 h time point, however, interphase cells showed minimal change, whereas mitotic cells already exhibited pronounced dispersal (Fig. 3c,d). These kinetics support a model in which ecDNA hub disruption begins during mitosis and subsequently emerges in interphase.

To test this model directly, we performed live-cell imaging using a locus-labelling strategy in which a 96× TetO array was engineered adjacent to the *MYC* gene encoded on ecDNA and visualised with TetR-mNeonGreen, while H2B-emiRFP670 marked chromosomal DNA (Fig. 3e; Extended Data Fig. 7). This approach enabled comparison of ecDNA organisation in the same cell before and after WEE1 degradation. ecDNA dispersal increased only in cells that progressed through mitosis during imaging (Fig. 3f,g), whereas cells that remained in interphase showed no obvious change (Fig. 3h). Consistently, direct analysis of mitotic cells revealed increased ecDNA dispersal after WEE1 degradation relative to DMSO-treated cells (Fig. 3i,j). Together, these findings support a model in which WEE1 degradation promotes ecDNA hub dispersal through mitotic progression, with mitosis serving as the point of initiation for subsequent daughter-cell ecDNA reorganisation.

## CDK1 and CDK2 mediate hub dispersal

Having established that WEE1d-induced hub dispersal originates during mitosis, we next investigated how WEE1 loss promotes ecDNA reorganisation at this stage. ecDNA remains associated with mitotic chromosomes^17^, raising the possibility that chromosome-associated DNA lesions might alter ecDNA organisation. In addition to restraining mitotic entry, WEE1 limits CDK activity during S phase and protects genome integrity by constraining replication initiation and replication stress^18^. We therefore asked whether WEE1 degradation generated DNA lesions that became enriched during mitosis.

TUNEL analysis at 6 h after WEE1 degradation revealed a marked increase in DNA damage in mitotic cells, whereas interphase cells showed little change relative to DMSO-treated controls (Fig. 4a,b). Mitotic TUNEL signal remained elevated at 24 h, when an increase was also detected in interphase cells (Fig. 4b). In WEE1d-treated cells, TUNEL intensity remained substantially below that of DNase I-treated positive controls. Together with the preserved nuclear morphology and lineage tracking showing that only 1 of 39 mitosis-entering cells died during mitosis, these findings indicate that the increased TUNEL signal was not dominated by terminal apoptotic fragmentation (Fig. 3b). Instead, they are consistent with DNA strand breaks becoming enriched or persisting during mitosis. Moreover, γH2AX was already elevated in interphase cells after 6 h of WEE1 degradation and was also elevated in mitotic cells, supporting the induction of DNA-damage signalling before mitotic entry (Extended Data Fig. 8a,b).

**Fig. 4.**
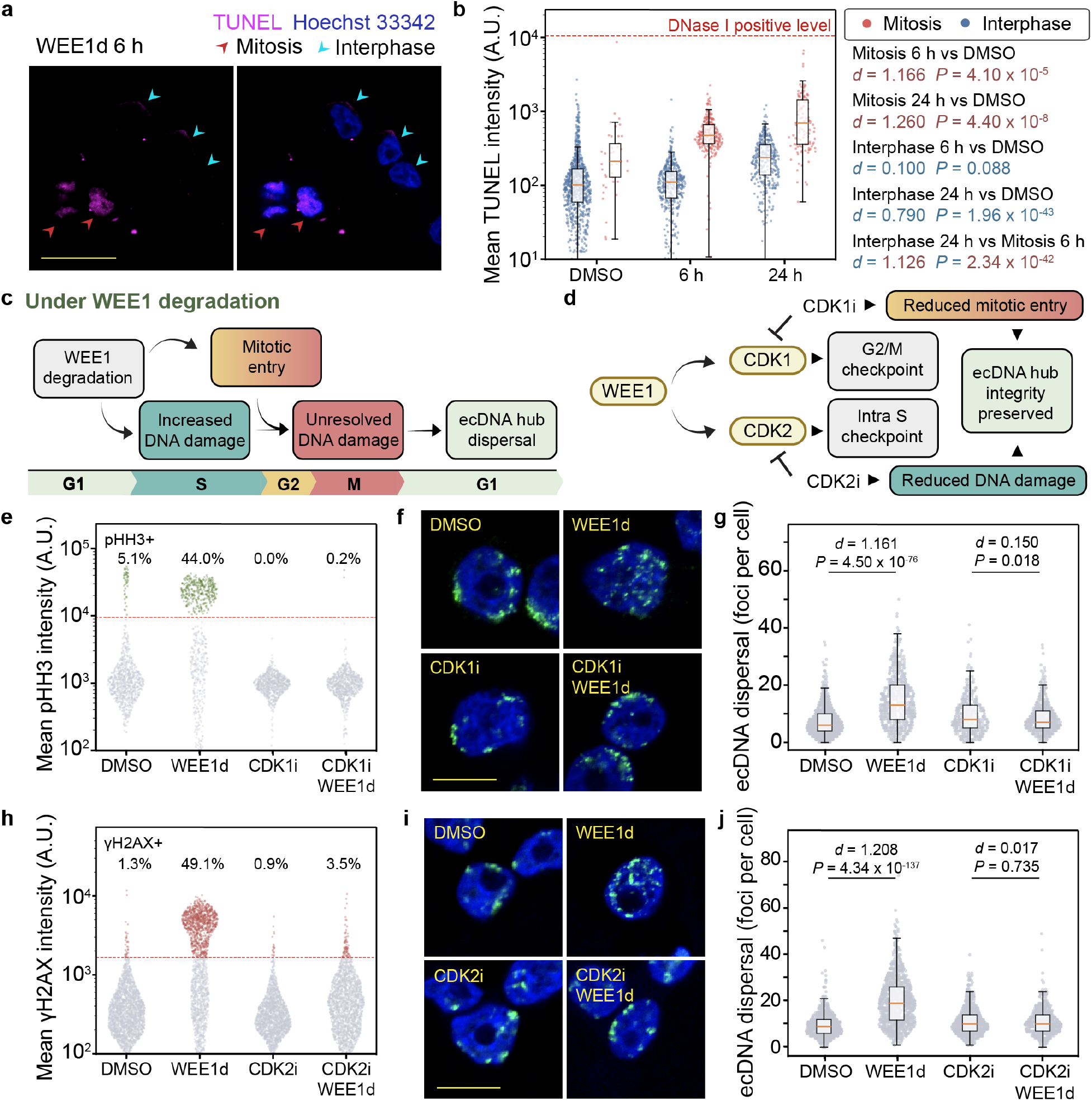
CDK1 and CDK2 mediate WEE1d-induced ecDNA hub dispersal. **a,b,** Representative TUNEL images of COLO320DM cells after 6 h of WEE1d treatment (**a**) and mean nuclear TUNEL intensity in interphase and mitotic cells treated with DMSO or WEE1d for 6 or 24 h (**b**; *n* = 2 biological replicates). Red and cyan arrowheads indicate mitotic and interphase cells, respectively. TUNEL is magenta and Hoechst 33342 is blue. The dashed red line in **b** indicates the mean TUNEL intensity of DNase I-treated positive-control cells. **c,d,** Models of WEE1d-induced ecDNA hub dispersal (**c**) and the predicted effects of CDK1 or CDK2 inhibition (**d**). **e–g,** Nuclear pHH3 intensity and pHH3-positive fraction after 6 h of treatment (**e**), and representative *MYC* DNA FISH images (**f**) and ecDNA dispersal, quantified as foci per cell (**g**), after 24 h of treatment with DMSO, WEE1d, CDK1i (RO-3306) or CDK1i plus WEE1d (*n* = 3 biological replicates). **h–j,** Nuclear γH2AX intensity and γH2AX-positive fraction after 6 h of treatment (**h**), and representative *MYC* DNA FISH images (**i**) and ecDNA dispersal (**j**) after 24 h of treatment with DMSO, WEE1d, CDK2i (BLU-222) or CDK2i plus WEE1d (*n* = 3 biological replicates). Points in **e,h** represent individual nuclei. Dashed lines denote positivity thresholds; percentages indicate positive cells. **a,f,i** show single-z-plane images. In **f,i**, *MYC* DNA FISH is green and Hoechst 33342 is blue. Concentrations: 2 μM WEE1d, 10 μM CDK1i and 125 nM CDK2i. Sample sizes are provided in Supplementary Table 1. Two-sided Welch’s *t*-tests were used in **b,g,j**, with absolute Cohen’s *d* reported. TUNEL intensities were log10-transformed for statistical testing. Box plots show medians and interquartile ranges; whiskers extend to 1.5× the interquartile range. Scale bars, 20 μm (**a**) and 10 μm (**f,i**).

These observations suggested a two-component model in which WEE1 degradation increases CDK activity across S phase and G2/M, promoting both DNA-damage accumulation and mitotic entry through CDK1- and CDK2-regulated programmes. The convergence of these effects could allow incompletely resolved lesions to persist into mitosis and promote the reorganisation of chromosome-associated ecDNA (Fig. 4c). This model predicts that perturbing either process should attenuate ecDNA hub dispersal (Fig. 4d).

To test this model, we used CDK1 and CDK2 inhibitors to preferentially perturb mitotic entry and damage accumulation, respectively. CDK1 inhibition with RO-3306 (CDK1i)^19^ markedly reduced the fraction of cells positive for phospho-histone H3 (pHH3), a marker of mitotic chromatin^20^, and restored ecDNA hub organisation under WEE1 degradation (Fig. 4e–g; Extended Data Fig. 8c). CDK2 inhibition with BLU-222 (CDK2i)^21^ suppressed WEE1d-induced γH2AX, a marker of DNA-damage signalling, and reduced, but did not abolish, the pHH3-positive population, while similarly restoring ecDNA hub organisation (Fig. 4h–j; Extended Data Fig. 8d,e). Together, these results support a model in which CDK1- and CDK2-regulated programmes make complementary contributions to WEE1d-induced ecDNA hub dispersal by jointly shaping DNA-damage accumulation and progression into mitosis.

## ecDNA hubs persist through interphase

Having established that WEE1d-induced ecDNA dispersal depends on mitotic passage, we next asked whether mitosis also shapes ecDNA hub organisation under unperturbed conditions. We first characterised the dynamics of ecDNA hubs during interphase. Using live-cell imaging, we tracked labelled ecDNA hubs larger than 2 μm² in cells with at least 12 h of trackable interphase, providing an extended window in which to assess hub persistence. In total, we followed 64 hubs in 50 cells over a 24-h acquisition period (Fig. 5a; Extended Data Fig. 9a). Unexpectedly, individual hubs could be linked across hourly frames on the basis of their approximate size and relative position within the nucleus (Extended Data Fig. 9a). Although hub boundaries and shapes changed over time, individual hubs often remained recognisable throughout interphase, suggesting that large-scale reorganisation during this period is limited. Consistently, the mean trackable duration of individual hubs was 19.1 h, corresponding to 87.2% of the mean trackable interphase duration per cell (21.9 h) (Fig. 5b). Thus, the large interphase ecDNA hubs analysed here were not rapidly appearing and disappearing assemblies, but instead exhibited substantial temporal persistence.

**Fig. 5.**
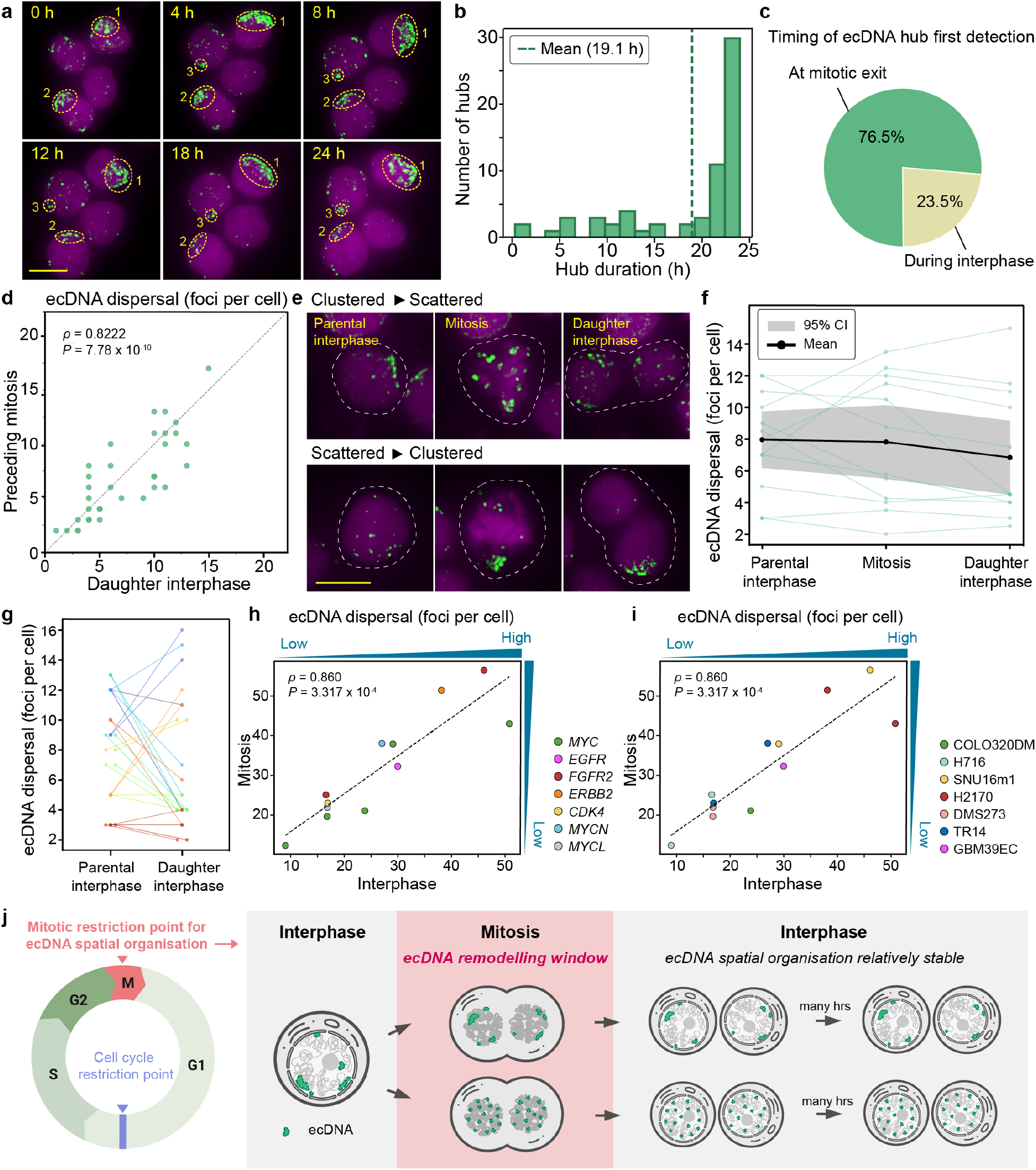
Interphase ecDNA hubs persist, whereas mitosis enables remodelling. **a– c,** Representative images of tracked ecDNA hubs (**a**), distribution of trackable hub durations (**b**) and timing of first hub detection (**c**) in DMSO-treated COLO320DM cells (*n* = 4 biological replicates). Hubs larger than 2 μm² were manually tracked in 24-h movies acquired hourly; only interphases longer than 12 h were retained. The dashed line in **b** marks the mean. Analysis in **c** was restricted to interphases beginning with a clearly observed mitotic exit; hubs were classified as first detected at mitotic exit or later during interphase. **d,** Correlation of the degree of ecDNA dispersal, quantified as foci per cell, between the last mitotic and first daughter-interphase states of matched lineages (*n* = 12 biological replicates). **e–g,** Representative clustered-to-scattered and scattered-to-clustered transitions (**e**), degree of ecDNA dispersal across parental interphase, mitosis and daughter interphase (**f**), and paired parental- and daughter-interphase measurements (**g**) (*n* = 5 biological replicates). **h,i,** Comparison of interphase and mitotic degree of ecDNA dispersal across seven ecDNA-positive cancer cell lines. Points represent cell line-oncogene pairs coloured by ecDNA-associated oncogene (**h**) or cell line (**i**); dashed lines denote *y* = *x*. **j,** Model of the mitotic restriction point, in which ecDNA configurations remodelled or maintained during mitosis are transmitted into daughter cells and remain comparatively stable during interphase. In **f**, coloured lines represent lineages, the black line denotes the mean and shading indicates the 95% confidence interval. Dashed white outlines in **e** mark tracked lineages; dashed yellow outlines in **a** highlight hubs. Images are SVCC-rendered maximum-intensity projections; ecDNA is green and chromosomal DNA is magenta. Sample sizes are provided in Supplementary Tables 1 and 2. Associations in **d,h,i** were assessed by two-sided Spearman correlation; **g** used a two-sided Wilcoxon test on lineage-level pairs (*P* = 0.4). Scale bars, 10 μm.

The extended trackability of ecDNA hubs suggested that, once established, they remain relatively stable during interphase. We therefore asked whether these hubs arise continuously during interphase or are primarily established around mitotic exit. To distinguish these possibilities, we first restricted the analysis to cells whose interphase trajectories began with a clearly observed mitotic exit, allowing hubs detected in the initial daughter-interphase frame to be distinguished from those first detected later. Of the hubs analysed, 76.5% were present in the initial daughter-interphase frame, whereas only 23.5% were first detected later during interphase (Fig. 5c). A broader analysis that also included trajectories already in interphase at movie onset yielded a similar distribution: 81.2% of hubs were present at movie onset or mitotic exit, whereas 18.8% were first detected later during interphase (Extended Data Fig. 9a). These results favour the possibility that interphase hubs are established around mitotic exit rather than continuously assembled during interphase, prompting us to ask whether daughter-interphase organisation reflects the preceding mitotic state. We therefore compared the last available mitotic frame of each cell with the first available daughter-interphase frame from the same lineage. Degree of ecDNA dispersal, quantified as the number of resolved foci per cell, was strongly correlated between the paired states, and individual large hubs could be visually followed across mitotic exit (Fig. 5d; Extended Data Fig. 9b). These observations indicate that the ecDNA dispersal state in daughter interphase closely reflects that present during the preceding mitosis.

## Mitosis remodels hubs across cancers

The observations above showed that interphase ecDNA hubs are relatively persistent and that daughter-interphase organisation is closely linked to the preceding mitotic state. Together, these findings raised the possibility that mitosis serves as a major remodelling window for ecDNA spatial organisation. To test this directly, we analysed cell lineages that progressed through mitosis and compared the degree of ecDNA dispersal across three sequential states: parental interphase, mitosis and daughter interphase. We quantified ecDNA dispersal as foci per cell, with higher values indicating more dispersed ecDNA. For the mitotic and daughter stages, measurements from the two separated chromosomal DNA masses in anaphase, telophase and daughter interphase were averaged to account for partitioning of duplicated ecDNA between the two daughter cells.

Individual lineages exhibited trajectories in which ecDNA became either more dispersed or more clustered during mitotic passage (Fig. 5e,f). At the population level, however, the mean degree of ecDNA dispersal remained relatively stable across parental interphase, mitosis and daughter interphase, indicating no consistent directional shift under baseline conditions (Fig. 5f). Thus, population averaging obscured substantial bidirectional changes within individual lineages. Direct comparison of paired parental- and daughter-interphase states likewise revealed marked lineage-specific changes, despite the absence of a population-level directional shift (Fig. 5g).

To assess the extent of parent-daughter similarity attributable to lineage identity, we compared the median lineage-averaged absolute parent-daughter difference with a null distribution generated by randomly reassigning daughter lineage identities. The observed difference was indistinguishable from the null distribution and coincided with its median (Extended Data Fig. 9c), indicating that matched daughter states were no more similar to their parental states than expected under random pairing for this metric. Considered together with the strong correspondence between mitotic and daughter-interphase states (Fig. 5d), these results support mitosis as a major window in which ecDNA hub organisation is remodelled before being transmitted into daughter interphase.

We next asked whether this principle extends across ecDNA-positive cancer models. Baseline ecDNA dispersal varies substantially among cancers, providing a broad dynamic range in which to test whether mitotic and interphase organisation remain coupled. If the mitotic state shapes subsequent interphase organisation, models with relatively compact interphase ecDNA should also exhibit compact mitotic ecDNA, whereas models with more dispersed interphase ecDNA should show correspondingly dispersed mitotic states.

To examine this, we analysed seven ecDNA-positive cell lines spanning colorectal, gastric and lung cancers, neuroblastoma and glioblastoma, and representing ecDNA-associated oncogenes including *MYC*, *FGFR2*, *ERBB2*, *MYCL*, *CDK4*, *MYCN* and *EGFR* (Fig. 5h,i; Extended Data Fig. 10a–e). To capture complementary aspects of ecDNA dispersal beyond foci number, we also quantified maximum cluster size and the fraction of total ecDNA signal contained within the largest cluster. All three metrics showed strong positive correlations between mitotic and interphase populations (Fig. 5h,i; Extended Data Fig. 10b–e). These relationships spanned models with different ecDNA-associated oncogenes and tumour origins, indicating that mitosis-interphase correspondence is not obviously dictated by either feature. Across this broad range of ecDNA dispersal states, the degree of dispersal in interphase consistently tracked with that in mitotic cells, supporting the generality of our finding that ecDNA organisation during mitosis is carried forward into interphase. An exploratory analysis further showed a positive, although weaker, correlation between mitotic and interphase populations in the intensity-weighted distance of ecDNA signal from the DNA-mass edge (Extended Data Fig. 10f,g). This finding suggests that mitosis-interphase correspondence may extend beyond hub clustering to radial ecDNA positioning. Together, these results are consistent with mitosis serving as a broadly shared remodelling window for ecDNA spatial organisation, with mitotic configurations carried forward into the subsequent interphase (Fig. 5j).

## Discussion

In this study, we combined imaging-based perturbation screening, fixed-cell analysis and live-cell tracking to investigate how ecDNA hubs are regulated across the cell cycle. Although ecDNA hubs have emerged as a prominent feature of ecDNA biology, the mechanisms governing their formation, disruption and persistence have remained poorly understood. Our results identify WEE1 as a regulator of ecDNA hub integrity and show that WEE1 loss disperses hubs in the subsequent interphase and reduces ecDNA-encoded oncogene expression. Rather than being continuously reconfigured throughout interphase, ecDNA hubs are remodelled during mitosis, and the resulting spatial states are frequently inherited and maintained throughout the subsequent interphase. Correspondence in ecDNA radial positioning between mitotic and interphase populations across cancer models further suggests that this cell-cycle-linked regulation may extend to broader features of ecDNA spatial organisation.

Together, these findings support a model conceptually analogous to, but distinct from, the classical cell-cycle restriction point. The discovery by Pardee and colleagues that passage through a G1 restriction point commits mammalian cells to complete the cell-division cycle was a seminal advance in cell biology^22,23^. The restriction point is now understood as a critical window during which cells integrate inputs from cell size, growth factor signalling, and metabolic state to commit to DNA replication and cell division, ultimately leading to the identification of key molecular regulators such as cyclins, Rb, and E2F proteins. By contrast, our findings suggest that the restriction point governing ecDNA spatial organisation, and by extension ecDNA function, occurs not in G1 but during mitosis (Fig. 5j). In this model, ecDNA spatial states generated or maintained during mitosis are preferentially propagated into daughter cells and remain constrained until the next mitotic remodelling window. Two features of this model warrant emphasis. First, it does not imply that ecDNA is static during interphase. Live-cell imaging revealed apparent local dynamics, including changes in hub size and shape, even as the broader organisational state remained comparatively constrained within a given cell cycle. Second, mitosis does not appear to enforce obligatory reorganisation in every cell. Under baseline conditions, some lineages changed little, whereas others became substantially more clustered or dispersed across mitotic passage. Mitosis therefore creates an opportunity for ecDNA reorganisation without imposing a uniform spatial outcome.

ecDNA associates with mitotic chromosomes through retention elements, resembling the chromosome-tethering strategies used by several episomal DNA viruses to retain their genomes through cell division^17^. This association raises the possibility that changes in the mitotic chromosome environment could directly reshape ecDNA organisation. The identification of WEE1 provided a mechanistic entry point into this relationship. WEE1 degradation induced DNA-damage signalling before mitosis and enriched DNA strand breaks during mitosis, consistent with complementary contributions from CDK1- and CDK2-regulated programmes. Although extensive crosstalk between CDK1 and CDK2 is likely to connect these downstream programmes, reducing the burden of mitotic DNA damage through either CDK1 or CDK2 inhibition restored ecDNA hub organisation, supporting a model in which the organisation of chromosome-tethered ecDNA is shaped by the broader state of mitotic chromosomes. In this model, WEE1 acts indirectly by generating an aberrant mitotic environment rather than functioning as a structural component of ecDNA hubs. Other perturbations that disrupt mitotic chromosome integrity may similarly affect ecDNA hubs, but produce weaker population-level phenotypes because mitotic cells constitute only a small fraction of an asynchronous population. WEE1 degradation may be particularly effective because it couples DNA damage with increased mitotic entry, enriching cells within this remodelling window.

Orderly mitosis helps re-establish nuclear architecture in daughter cells. Features such as chromosome territories and centromere positioning can persist across cell cycles while also being reorganised during mitosis^24,25^. Higher-order ecDNA organisation may represent another feature shaped during this transition. A complementary possibility is that changes in mitotic kinase activity could remodel hub-associated factors and weaken the intermolecular interactions that support hub integrity^26^. Mitosis drives extensive biochemical remodelling of nuclear compartments, including nucleoli, nuclear speckles and PML bodies, and DYRK3-dependent phosphorylation has been shown to promote the dissolution of several membraneless organelles^27–30^. These precedents suggest that mitotic phosphorylation may similarly alter the molecular interactions that contribute to ecDNA hub organisation. Defining how ecDNA engages mitotic chromosomes, whether and how hub-associated factors are remodelled and how these processes shape daughter-cell organisation will be important directions for future studies.

Our results revise expectations for ecDNA behaviour during interphase. Because ecDNAs lack centromeres and are not embedded within linear chromosomes, they might be expected to move broadly throughout the nuclear space. Instead, large ecDNA hubs were locally dynamic but temporally persistent, suggesting that ecDNA organisation is more constrained within a given interphase. Spatial diversity among ecDNAs may therefore emerge through the collective behaviour of many copies, periodic reconfiguration during mitosis and variation among individual cells, rather than through unrestricted long-range movement during interphase. Persistent cell-to-cell differences in ecDNA organisation could consequently contribute a spatial dimension to intratumoural heterogeneity by shaping the regulatory interactions and transcriptional states available to individual cancer cells.

Finally, these findings raise the possibility that mitosis represents a therapeutically accessible window for perturbing ecDNA-dependent oncogenic states. Altering mitotic progression or the molecular interactions that connect ecDNA to mitotic chromosomes could reshape ecDNA organisation and its associated transcriptional programmes. Establishing which ecDNA configurations support cancer-cell fitness, and whether these states can be selectively disrupted without broadly increasing genome instability, will be important for translating the mitotic restriction-point model into therapeutic strategies.

## Methods

### Cell culture

Parental COLO320DM, SNU16, H2170 and H716 cells were obtained from ATCC, and DMS273 cells from Sigma-Aldrich. SMS-KAN cells were provided by the laboratory of Anton G. Henssen. The monoclonal SNU16m1 line was derived from parental SNU16 cells by single-cell cloning following lentiviral expression of dCas9-KRAB, as described previously^6^. GBM39EC is a patient-derived glioblastoma neurosphere line established previously^7,33^, and TR14 neuroblastoma cells were provided by J. J. Molenaar^9^. COLO320DM-TG19, which carries a TetO array adjacent to *MYC*-containing ecDNA and stably expresses TetR-mNeonGreen and H2B-emiRFP670, was generated previously^9^. COLO320DM-FUCCI was generated as described below. Cell-line identifiers and complete culture conditions are provided in Supplementary Table 3.

SNU16m1 cells were maintained in DMEM/F12 with 10% FBS and 1% PSG. COLO320DM and its derivatives were maintained in DMEM with 10% FBS and 1% PSG. GBM39EC cells were maintained in DMEM/F12 supplemented with B-27, PS, GlutaMAX, EGF, FGF and heparin. H716, H2170, DMS273 and SMS-KAN cells were maintained in ATCC-formulated RPMI 1640 with 10% FBS and 1% PSG, and TR14 cells in RPMI 1640 with 20% FBS and 1% PS. Reagent suppliers and identifiers are provided in Supplementary Table 9. Cells were cultured at 37 °C with 5% CO₂ and periodically tested for mycoplasma contamination, primarily using the MycoAlert PLUS assay; all tested negative.

### Generation of COLO320DM-FUCCI H2B-emiRFP670 cells

Parental COLO320DM cells were cotransfected in six-well plates with 1.6 µg tFucci(CA)5 reporter plasmid and 0.4 µg Super PiggyBac transposase vector using 2 µl X-tremeGENE HP in 200 µl Opti-MEM per well. Complexes were incubated for 15 min at room temperature, added to cells overnight and replaced with fresh medium the following day. Cells were expanded without enrichment.

H2B-emiRFP670 lentivirus was produced in HEK293T cells by transfecting 900 ng psPAX2, 100 ng pMD2.G and 1 µg pHR-H2B-emiRFP670-GB1-dWPRE using 10 µl TransIT-LT1 in 250 µl Opti-MEM. Viral supernatant was collected after approximately 72 h and passed through a 0.45-µm filter. FUCCI(CA)-expressing COLO320DM cells were transduced with 250 µl viral supernatant in medium containing 10 µg ml⁻¹ polybrene and 20 mM HEPES in a final volume of 2 ml, followed by centrifugation at 800g for 2 h at room temperature. The viral medium was then replaced. After expansion, cells were sorted using all three fluorescent channels, and the pooled population was retained to obtain relatively homogeneous reporter expression. Commercial reagents are listed in Supplementary Table 9.

### Chemicals

Cells were treated with DMSO or the indicated compounds for the durations specified in the figure legends, with the final DMSO concentration matched across conditions. A focused nuclear protein-targeting compound library containing ZNL-02-096, referred to as WEE1d, was provided by M. M. Hassan and Nathanael S. Gray. Compounds were prepared as 10 mM stocks in DMSO. Library compounds were used at 10 µM in the hiFISH-96 screen. Unless otherwise indicated, WEE1d, MK-1775 (WEE1i), BLU-222 (CDK2i), MG132 and RO-3306 (CDK1i) were used at 2 µM, 500 nM, 125 nM, 10 µM and 10 µM, respectively. Cells were pretreated with MG132 for 2 h before WEE1d addition, whereas CDK1i or CDK2i was added concurrently with WEE1d. Compound sources are provided in Supplementary Table 9.

### Imaging-based compound screen

The hiFISH-96 screen was performed in COLO320DM cells. Cells grown to 70–80% confluency were seeded at 80,000 cells per well in coated 96-well imaging plates (Supplementary Table 4) and treated 5 h later. The screening plate contained 43 compound-treated and five DMSO-control wells; compound identities are provided in Supplementary Data 1. For repeated-control experiments, 60 DMSO-treated wells were processed, of which 54 passed the quality-control criteria described below. At 24 h after treatment, 16% PFA was added directly to the medium to a final concentration of 4%, and cells were fixed for 10 min at room temperature before hiFISH-96 detection of *MYC*-containing ecDNA. Four compounds causing overt cell loss were excluded before imaging. The remaining 39 compounds were evaluated using ecDNA dispersal, quantified as the mean number of spatially resolved ecDNA foci per cell, as the primary screening metric and ranked relative to pooled DMSO controls. ecDNA radial distribution was analysed as a secondary readout. Imaging and quantitative analysis were performed as described below.

### DNA fluorescence in situ hybridisation

DNA FISH was performed using heat-denaturation, acid-denaturation, metaphase-spread or hiFISH-96 workflows according to sample format and assay requirements (Supplementary Table 5). Probe information is provided in Supplementary Table 6 and reagent information in Supplementary Table 9. Probes were diluted in the hybridisation buffer supplied with the commercial probes.

For coverslip-based FISH, cells were fixed with 4% PFA for 10 min at room temperature. For heat denaturation, cells were permeabilised with 0.5% Triton X-100 in PBS for 15 min, dehydrated through 70%, 85% and 100% ethanol and incubated with probes diluted 1:10. Samples were sealed with rubber cement, denatured at 80 °C for 10 min and hybridised overnight at 37 °C. For acid denaturation, samples were incubated with 0.7% Triton X-100 and 0.1 M HCl in PBS for 10 min on ice, followed by 1.9 M HCl for 30 min at room temperature and ethanol dehydration. Probes were diluted 1:5, pre-denatured at 75 °C for 3 min and hybridised overnight at 37 °C. When combined with immunofluorescence, immunostaining preceded acid-denaturation FISH. After hybridisation, coverslips were briefly rinsed with 0.4× SSC and washed twice for 15 min with 2× SSC containing 0.1% Tween-20 at room temperature. DNA was stained with Hoechst 33342, and samples were mounted with ProLong Gold, cured for approximately 24 h and stored at −20 °C until imaging.

For metaphase-spread FISH, air-dried slides were dehydrated through 70%, 85% and 100% ethanol for 2 min each. *MYC* and TetO probes were diluted 1:14 and 1:35, respectively, applied beneath coverslips, denatured at 75 °C for 3 min and hybridised overnight at 37 °C in a humidified chamber. Slides were washed sequentially with 0.4× SSC, 2× SSCT and 2× SSC for 2 min each, stained with 100 ng ml⁻¹ DAPI, washed, air-dried and mounted with ProLong Diamond.

### hiFISH-96 plate-format DNA FISH

hiFISH-96 was used for the compound screen and other fixed-cell DNA FISH experiments performed in 96-well imaging plates. Plates were coated as described in Supplementary Table 4, and cells were seeded and treated as indicated. To minimise cell loss, 16% PFA was added directly to the medium to a final concentration of 4%, and cells were fixed for 10 min at room temperature.

hiFISH-96 was developed to enable high-throughput DNA FISH in a 96-well format while minimising probe consumption. Published multi-well DNA FISH workflows typically use 10–15 μl of probe mixture per well in 384-well plates^34^. Scaling this volume proportionally to the larger well area of a 96-well plate would require approximately 40–60 μl per well, limiting cost-effective high-throughput implementation. hiFISH-96 instead uses 3.5 μl of probe mixture per well, representing an approximately 11-to 17-fold reduction relative to the area-scaled volume while permitting analysis of larger cell populations. This increased sampling capacity is particularly useful for ecDNA analyses, in which cell-to-cell heterogeneity is substantial.

The heat-denaturation DNA FISH workflow was adapted to the reduced-volume plate format as follows. Fixed cells were permeabilised with 0.5% Triton X-100 in PBS for 15 min. Ethanol dehydration was generally omitted because residual liquid could be removed efficiently from the wells. FISH probes were diluted 1:10 in hybridisation buffer and applied at 3.5 µl per well. Each sample well was covered with a 7-mm glass coverslip and sealed with rubber cement to prevent evaporation. Rubber cement was also applied within neighbouring unused wells to reinforce attachment of the glass bottom to the plate frame during heating and washing. Samples were denatured at 80 °C for 10 min and hybridised overnight at 37 °C in a humidified dark chamber.

After hybridisation, rubber cement and coverslips were removed from sample wells while the supporting cement in neighbouring wells was retained. Samples were briefly rinsed with 0.4× SSC, washed twice for 15 min with 2× SSCT and stained with Hoechst 33342. The supporting cement was then removed, and the plate was dried. Each well was mounted with 3.5 µl ProLong Gold beneath a new 7-mm coverslip, cured for approximately 24 h and stored at −20 °C until imaging. Reagents and materials are listed in Supplementary Table 9.

### Metaphase spread

COLO320DM-TG19 cells grown to 70–80% confluency in 6-cm dishes were treated with 0.1 μg ml⁻¹ KaryoMAX Colcemid for 4 h. Mitotic cells were collected by shake-off, and the remaining adherent cells were trypsinised for 2 min. The two fractions were combined and centrifuged at 400g for 4 min. Cell pellets were resuspended in 600 µl of 75 mM KCl and incubated at 37 °C for 20 min. An equal volume of Carnoy’s fixative (methanol:acetic acid, 3:1) was added dropwise, followed by centrifugation at 950 g for 2 min. Pellets were washed three times with 600 µl Carnoy’s fixative and resuspended in 200 µl. Ten microlitres of cell suspension was dropped from approximately 15 cm onto glass slides pre-warmed and humidified using warm, moist air, and allowed to air-dry. Reagents are listed in Supplementary Table 9.

### Immunofluorescence

COLO320DM cells were seeded into coated 96-well imaging plates (Supplementary Table 4) and treated for 6 h with DMSO, WEE1d, CDK1i or CDK2i alone or in combination with WEE1d. To minimise cell loss, 16% PFA was added directly to the medium to a final concentration of 4%, and cells were fixed for 10 min at room temperature.

Cells were permeabilised with 0.5% Triton X-100 in PBS for 15 min and blocked with 3% BSA in PBS containing 0.1% Tween-20 (PBST) for at least 1 h. Samples were incubated overnight at 4 °C with rabbit anti-pHH3, mouse anti-γH2AX or both at 1:300. After three PBST washes, appropriate fluorophore-conjugated secondary antibodies were applied at 1:500 for 30 min at room temperature. Samples were subsequently protected from light, washed twice with PBST, stained with Hoechst 33342 for 30 min and imaged in PBS. Antibody and reagent information is provided in Supplementary Tables 7 and 9.

Z-stack images were acquired as detailed in Supplementary Table 8. Mean nuclear pHH3 and γH2AX intensities and the corresponding positive-cell fractions were quantified. The pHH3 threshold was selected using DMSO-treated cells to distinguish morphologically verified mitotic and interphase nuclei. The γH2AX threshold was selected by comparing DMSO- and WEE1d-treated intensity distributions, with DMSO defining the background distribution. Each threshold was then applied unchanged across all conditions within the corresponding experiment.

### TUNEL assay

TUNEL staining was performed using the Click-iT TUNEL Alexa Fluor Imaging Assay according to the manufacturer’s instructions. COLO320DM cells were seeded onto coated coverslips (Supplementary Table 4), treated with DMSO or WEE1d for 6 or 24 h and fixed with 4% PFA for 10 min at room temperature. Cells were permeabilised with 0.25% Triton X-100 in PBS for 20 min and washed twice with deionised water. For the positive control, fixed and permeabilised untreated cells were incubated with the DNase I solution supplied with the kit for 30 min at room temperature.

Samples were incubated in TdT reaction buffer for 10 min at 37 °C, followed by TdT reaction mixture for 60 min at 37 °C. After two washes with 3% BSA in PBS, samples were incubated with Click-iT reaction cocktail for 30 min at 37 °C protected from light, washed, stained with Hoechst 33342 and mounted with ProLong Gold. Reagents are listed in Supplementary Table 9.

Z-stack images were acquired as detailed in Supplementary Table 8. For each nucleus, the plane with the highest mean Hoechst 33342 intensity was selected, and mean TUNEL intensity was quantified in that plane as a measure of DNA strand breaks.

### Live-cell imaging and image visualisation

COLO320DM-TG19 and COLO320DM-FUCCI cells were seeded into poly-D-lysine-coated 96-well imaging plates (Supplementary Table 4) one day before imaging. The environmental chamber was equilibrated to 37 °C and 5% CO₂ under humidified conditions. Immediately before acquisition, DMSO or WEE1d was diluted in complete medium containing ProLong Live Antifade and added directly to the existing medium with gentle mixing, yielding final concentrations of 2 µM WEE1d and 1:100 ProLong Live Antifade. DMSO controls received an equivalent DMSO volume. Acquisition settings are provided in Supplementary Table 8 and reagents in Supplementary Table 9. Images displayed in figures were processed using the Leica LAS X Small Volume Computational Clearing method; all quantitative analyses used raw images.

### Image acquisition

Fluorescence channels were selected according to the fluorophores or stains used in each experiment. Illumination intensity and exposure time were optimised for each experiment and held constant across samples included in direct comparisons. Microscopes, objectives, acquisition software, spatial sampling and experiment-specific z-stack and time-lapse settings are summarised in Supplementary Table 8.

### Image preprocessing and feature segmentation

The analysis workflow is summarised in Extended Data Fig. 1a. Analyses used custom Python scripts, Cellpose v.3.1.1.1^35^, ilastik v.1.4.0^36^ and napari. Automated procedures were applied uniformly across directly compared conditions, and sample identities were masked during manual curation. Exact implementation parameters are provided in the deposited code.

#### Image preprocessing and registration

Leica TCS SP8 and Leica DMi8 images were analysed directly. Nikon spinning-disk images were flat-field corrected using normalised and smoothed maximum-intensity projections sampled across each acquisition batch. When chromatic aberration was evident, corresponding points were manually identified in at least five fields sampled across conditions and pooled to calculate one affine transformation per batch, using the nuclear channel as reference. The resulting transformation was applied uniformly to all images in the batch before segmentation and feature quantification.

#### Nuclear segmentation

Nuclear or DNA-mass regions were segmented from Hoechst 33342, DAPI or H2B-emiRFP670 signal using Cellpose. The default nuclei model was used for Leica TCS SP8 datasets, including the screen, repeated DMSO controls and dose-response experiments. For other datasets, one custom model was trained per cell type using manually annotated masks. Fixed-cell SP8 images were segmented as single planes; other fixed-cell z-stacks were segmented as both individual planes and maximum-intensity projections; and live-cell images were segmented as maximum-intensity projections.

#### Nuclear-mask filtering and quality control

For live-cell datasets, nuclear or DNA-mass masks were manually corrected using custom Python interfaces in napari when required. For the screen and repeated DMSO controls, nuclear masks were filtered using predefined morphological criteria and evaluated by masked inspection of randomly selected fields until approximately 100 nuclei had been reviewed per sample or all fields had been examined. Samples with less than 90% initial segmentation accuracy were manually corrected and retained only if they passed predefined post-correction segmentation and FISH-intensity criteria. Overall, 54 of 60 repeated DMSO-control wells passed these criteria.

Other interphase-only fixed-cell datasets were filtered using batch-specific random-forest classifiers trained on manually curated fields sampled across conditions. The classifiers excluded image-border objects, debris, mitotic cells and fused or abnormally large masks using morphological, intensity and local-crowding features. A related classifier retained masks and distinguished interphase from mitotic DNA masses in the TUNEL analysis. Fixed-cell DNA FISH analyses requiring separate interphase and mitotic measurements were instead curated manually because mitotic cells were rare. For z-stack datasets, curated maximum-projection masks defined retained nuclei or DNA masses, and individual-plane masks were matched using overlap, area and centroid position.

#### ecDNA segmentation and object filtering

For the screen and repeated DMSO controls, FISH-intensity thresholds were selected separately for each field to yield a positive non-zero pixel fraction closest to 9.295%. This target was empirically established from repeated DMSO controls based on segmentation consistency and visual agreement and applied unchanged to the screen, which used the same cell type and acquisition settings.

Other fixed-cell FISH datasets were segmented using batch-specific ilastik classifiers trained to distinguish FISH signal from background. Fields sampled across all conditions contributed to training or validation, and one model was applied across each batch. Live-cell ecDNA was segmented from maximum-intensity projections using one model per movie, with manual correction when necessary. Connected objects smaller than approximately 0.07 µm² in SP8 datasets or 0.035 µm² in Nikon and DMi8 datasets were excluded. Parameters were held constant across directly compared conditions.

### ecDNA feature quantification

#### ecDNA clustering analysis

Segmentation was performed on raw images or, when applicable, flat-field-corrected and channel-registered images. Intensity measurements used background-corrected images. Background was estimated for each field from randomly sampled non-overlapping regions below the Otsu threshold in all selected channels, excluding manually defined regions. For maximum-intensity projections, the field-level mean background was subtracted from the image. For z-stacks, the same sampling mask was applied to each plane to estimate and subtract plane-specific background. Negative values were set to zero.

ecDNA dispersal was used as the primary metric and was defined as the number of segmented ecDNA objects within the corresponding nuclear or mitotic DNA-mass region. For most fixed-cell DNA FISH datasets, this metric and other ecDNA features were extracted from an acquired single-z-plane image or a plane selected separately for each nucleus from a z-stack. For analyses restricted to interphase cells, a central nuclear plane was preferred because it more clearly resolved ecDNA clustering. A reference plane was first identified from the maximum DNA-stain signal within each nucleus. Because this commonly represented the basal nuclear surface, the analysis plane was selected at least 0.5 µm above it, towards the nuclear centre; for datasets acquired at 1-µm intervals, the next plane 1 µm above the reference was used. For comparisons including interphase and mitotic cells, the plane with the greatest integrated ecDNA intensity across segmented ecDNA objects within each nuclear or mitotic DNA-mass mask was selected to improve signal recovery in rare mitotic cells while enabling consistent comparison between cell-cycle states. Curated maximum-projection masks defined nuclei or DNA masses eligible for analysis, and cells without a corresponding valid single-plane mask were excluded.

Live-cell ecDNA features were measured from maximum-intensity projections to improve recovery of TetR-mNeonGreen-labelled ecDNA. For the cross-cell-line analyses in Fig. 5h,i and Extended Data Fig. 10b–e, we additionally quantified maximum ecDNA cluster size, defined as the area of the largest segmented object, and the fraction of ecDNA signal within the largest cluster, calculated as its integrated background-corrected intensity divided by the summed intensity of all segmented ecDNA objects in the cell. These features were measured from maximum-intensity projections to capture overall ecDNA organisation and large hub structures. Radial-positioning measurements in Extended Data Fig. 10f,g instead used single-z-plane images to avoid projection-induced distortion. Each cross-cell-line value represents the population mean for one cell line-oncogene pair and cell-cycle state. Previously generated H716 and TR14 DNA FISH datasets^9^ were reanalysed using the same workflow.

#### ecDNA radial-positioning analysis

Radial positioning was quantified using population-level radial enrichment profiles or per-cell intensity-weighted distance to the DNA-mass edge. For radial enrichment, ecDNA-positive pixels were aggregated across nuclei according to their distance from the nuclear edge. Enrichment in each distance bin was calculated as the fraction of all FISH-positive pixels in that bin divided by the corresponding fraction of nuclear pixels. Curves were smoothed by cubic-spline interpolation. Peak location was the position of maximum enrichment, and right 75% width was the distance from the peak to the position at which enrichment decreased to 75% of the peak on the side further from the nuclear edge.

For per-cell analysis, the distance from the closest pixel of each segmented ecDNA object to the nearest DNA-mass edge was measured. Object distances were averaged within each cell, weighted by integrated background-corrected object intensity, and population-level values were calculated as the mean of these per-cell measurements.

### Interphase hub persistence analysis

Live-cell movies were screened for cells with more than 12 h of trackable interphase. ecDNA hubs larger than 2 μm² were manually tracked forwards and backwards across consecutive hourly frames using their position, size, shape and signal continuity. Trackable duration was defined as the total period over which a hub could be followed. If a hub appeared to disappear transiently, that frame was recorded as undetected and tracking continued only if the hub could be reassigned within the next two frames; otherwise, the track was terminated.

Total analysable interphase duration was the period over which the corresponding cell could be followed continuously during interphase. The population-level proportion of trackable interphase was calculated as the mean hub-track duration divided by the mean analysable interphase duration per cell. Hubs were classified as present at movie onset, first detected in the initial daughter-interphase frame following mitosis or first detected later during interphase.

### Live-cell lineage analysis

Individual nuclei were manually tracked using their size, position and H2B-emiRFP670 signal. Mitotic frames were identified from chromosomal DNA morphology, with chromosome condensation used as the primary criterion.

For event-level analyses, separated DNA masses were measured individually. For comparisons of mitotic ecDNA dispersal between DMSO- and WEE1d-treated cells (Fig. 3j), the last available mitotic frame from each lineage was used; frames containing one or two separated DNA masses contributed one or two measurements, respectively. For mitosis-to-daughter comparisons (Fig. 5d), each DNA mass in the last mitotic frame was paired by spatial correspondence with the appropriate nucleus in the first daughter-interphase frame. For parental-to-daughter comparisons (Fig. 5g and Extended Data Fig. 9c), the last parental-interphase frame was compared with the first daughter-interphase frame. Each daughter formed one parent-daughter pair, and pairs from the same lineage were identified accordingly. Daughter measurements were averaged within each lineage for statistical testing in Fig. 5g.

For lineage-level analyses, measurements from separated DNA masses were averaged within each frame. This rule was applied to comparisons between the beginning and end of imaging (Fig. 3g,h) and across parental interphase, mitosis and daughter interphase (Fig. 5f). For Fig. 3g,h, ecDNA dispersal was compared between the 0-h and 24-h frames. For Fig. 5f, parental-interphase values were averaged across the final two frames before mitotic entry, mitotic values across all mitotic frames and daughter-interphase values were taken from the first available daughter-interphase frame.

For the permutation analysis in Extended Data Fig. 9c, the observed median lineage-averaged absolute parent-daughter difference was compared with a null distribution generated by randomly reassigning daughter identities among lineages. The median lineage-level distance was recalculated for each of 10,000 permutations using a random seed of 12,345. A lower-tail permutation test assessed whether matched parent-daughter families were more similar than randomly paired families. The permutation *P* value was calculated as the fraction of null statistics less than or equal to the observed statistic, with a plus-one correction.

### FUCCI(CA) cell-cycle analysis

FUCCI(CA) live-cell movies were segmented as described above and tracked using ultrack v.0.7.1^37^. Tracks were manually verified and corrected to ensure tracking accuracy, including correction of tracking nodes and links. Mitotic frames were manually curated based on chromosomal DNA morphology as described above.

Mean fluorescence intensities in the hGem-h2-3 and hCdt1-AzaleaB5 channels were measured using locally dilated masks centred on the tracking nodes. G1 exit was manually identified from the resulting intensity traces. A decrease in hCdt1-AzaleaB5 signal, corresponding to hCdt1 loss, was required to define G1 exit and was supported by an accompanying increase in hGem-h2-3 signal when present^15,38^.

M-phase duration was measured only from tracks containing a complete M phase; tracks that began in M phase or remained in M phase at the final frame were excluded. The G1-exit-to-M interval was defined as the time from FUCCI-defined G1 exit to the first mitotic frame and was measured only from tracks containing both events.

For the 24-h cell-fate analysis in Fig. 3b, 50 cells that were in interphase at movie onset were randomly selected across two biological replicates and manually tracked. Cells were classified according to whether they entered mitosis during imaging and, for mitosis-entering cells, whether they completed mitosis, remained arrested in mitosis at the end of imaging or died during mitosis. Post-mitotic cells were further classified as alive and trackable, dead or lost to tracking. For the 24-h endpoint analysis, all trackable cells in interphase at the final time point were counted individually and classified according to whether they had passed through mitosis during imaging. Cells that were dead, in mitosis or no longer trackable at 24 h were excluded from this endpoint analysis.

### Hi-C library preparation

Hi-C libraries were generated using a modified HiChIP workflow^39^, with the chromatin immunoprecipitation step omitted. Buffers and procedures not otherwise specified followed the published protocol; reagents are listed in Supplementary Table 9. COLO320DM cells were treated with DMSO or WEE1d for 24 h. Two biological replicates containing 10 million cells per condition were processed in aliquots of 1–2 million cells. Cells were crosslinked with freshly prepared 2% methanol-free formaldehyde for 10 min at room temperature with rotation, quenched with 125 mM glycine on ice, washed and pelleted.

Crosslinked cells were lysed, and isolated nuclei were permeabilised with SDS and digested with 150 U DpnII per 10 million cells for 2 h at 37 °C. Restriction-fragment overhangs were filled in and biotinylated, followed by overnight proximity ligation with T4 DNA ligase at room temperature. Nuclei were lysed and sonicated using a Covaris instrument (fill level 10, duty cycle 5, peak incident power 140, 200 cycles per burst and 4 min). Clarified lysates were treated directly with Proteinase K for 45 min at 55 °C and reverse-crosslinked for 1.5 h at 67 °C before DNA purification.

Biotinylated ligation products were captured using Streptavidin C-1 beads and tagmented with adapter-loaded Tn5 transposase for 10 min at 55 °C. Libraries were amplified directly from the beads using dual-indexed Nextera-compatible primers and size-selected with 0.8× SPRISelect beads to retain fragments larger than 300 bp. Libraries were quantified by Qubit and Bioanalyzer and sequenced on an Illumina NovaSeq X Plus using 2 × 150-bp reads.

### Hi-C analysis

Hi-C paired-end sequencing reads were processed using HiC-Pro v.3.1.0^40^. Reads were aligned to the hg38 reference genome, filtered for valid interaction pairs, deduplicated and assigned to GATC restriction fragments using default HiC-Pro settings unless otherwise indicated. Genome-wide contact matrices were generated at multiple resolutions. Contact maps shown in Fig. 2e were visualised using Juicebox.

For ecDNA contact-redistribution analysis, raw 25-kb HiC-Pro contact matrices were normalised by total contact count using counts-per-million normalisation. Bins overlapping the COLO320DM ecDNA amplicon were defined using its genomic coordinates. ecDNA self-contact was calculated as the mean off-diagonal contact among amplicon-overlapping bins. External contact was calculated as the mean contact between amplicon-overlapping bins and all non-amplicon genomic bins. For each biological replicate, WEE1d-treated values were divided by the corresponding DMSO-treated values to calculate fold changes in ecDNA self-contact and external contact. Values from two biological replicates are shown in Fig. 2f.

### RNA-sequencing library preparation

COLO320DM cells were treated for 24 h with DMSO, WEE1d or WEE1i, with three biological replicates per condition. Total RNA was isolated from 1–2 million cells per sample using the RNeasy Plus Mini Kit. Stranded total-RNA libraries were prepared from 100 ng RNA using the TruSeq Stranded Total RNA Library Prep Kit with Illumina Ribo-Zero Plus rRNA depletion, according to the manufacturer’s protocol. Kits and reagents are listed in Supplementary Table 9. Libraries were sequenced on an Illumina NextSeq 550 system to generate 2 × 75-bp paired-end reads.

### RNA-sequencing analysis

Demultiplexed, ribosomal RNA-depleted paired-end reads were aligned directly to the hg38 human reference genome using STAR v.2.7.10b^41^. The genome index was generated using GENCODE v44 annotations^42^ and sjdbOverhang = 74. Uniquely mapped reads were retained using a mapping-quality threshold of 255.

Exon- and intron-associated fragments were quantified separately using featureCounts v.2.0.1^43^ in paired-end, reverse-stranded mode (-p -B -C -s 2). Exonic fragments were assigned using the GENCODE v44 exon annotation. Gene-level intronic regions were defined by subtracting the genome-wide union of annotated exons from each gene body. Intronic abundance was used as a proxy for nascent transcription, whereas exonic abundance represented predominantly steady-state RNA. Exon and intron count matrices were analysed separately using DESeq2 v.1.42.1^44^. WEE1d and WEE1i were each compared with DMSO using Wald tests, with *P* values adjusted using the Benjamini-Hochberg procedure.

Preranked gene-set enrichment analysis was performed separately for each comparison and RNA feature class using GSEApy v.1.3.1^45,46^. Genes were ranked by the DESeq2 Wald statistic. When multiple Ensembl identifiers mapped to one gene symbol, the entry with the largest absolute statistic was retained. Ranked lists were tested against the MSigDB Hallmark 2020 collection^47^ using 10,000 permutations, a random seed of 42 and gene-set sizes of 10–500. HALLMARK_MYC_TARGETS_V1 and HALLMARK_MYC_TARGETS_V2 were analysed separately.

### Immunoblotting

Cells were seeded in six-well plates at approximately 60–70% confluence and treated with DMSO or the indicated concentrations of WEE1d for 24 h. Cells were washed, detached with TrypLE and lysed in RIPA buffer supplemented with protease and phosphatase inhibitors. Lysates were sonicated using a Bioruptor, clarified by centrifugation at 16,000g for 20 min at 4 °C and quantified using a BCA assay.

Equal amounts of protein were denatured in LDS sample buffer containing 5% β-mercaptoethanol, separated on NuPAGE Bis-Tris gels and transferred to nitrocellulose membranes using an iBlot 2 system. Membranes were blocked with 3% BSA in PBST for at least 1 h at room temperature and incubated overnight at 4 °C with antibodies against WEE1 or GAPDH at dilutions of 1:100 and 1:1,000, respectively. After washing, membranes were incubated with IRDye secondary antibodies at 1:15,000 for 1 h at room temperature and imaged using a LI-COR imaging system. Antibodies are listed in Supplementary Table 7, and other reagents are listed in Supplementary Table 9.

### Statistical analysis

Statistical analyses were performed using Python unless otherwise stated. Statistical tests, sample sizes, biological replicates and definitions of *n* are provided in the figure legends and Supplementary Tables 1 and 2. Independent experiments used separate cell-culture preparations and treatments. Individual observations represent cells, ecDNA hubs, cell pairs or lineages, as indicated. For fixed-cell imaging analyses displayed as representative experiments, measurements from one biological replicate are shown to preserve the underlying single-cell distribution. Biological replicates were analysed separately and showed consistent treatment effects. Data were not pooled because batch-specific differences in imaging and segmentation could affect absolute feature values, whereas treatment and control conditions were acquired and analysed together within each experiment.

Tests were two-sided unless otherwise specified. No formal tests of normality were performed. Multiple-testing corrections were applied only where indicated. Welch’s *t*-test was used for two independent groups, Wilcoxon signed-rank tests for paired comparisons and Spearman’s rank correlation for associations between continuous variables. Parent-daughter similarity was assessed using the permutation analysis described under Live-cell lineage analysis. Cohen’s *d* was calculated as the difference between group means divided by the pooled standard deviation and is reported as an absolute value. For TUNEL analyses, Cohen’s *d* was calculated using log_10_-transformed intensities.

For the compound screen, up to 100 nuclei from each treatment were randomly sampled and compared with an equal number from the pooled DMSO controls using two-sample Kolmogorov-Smirnov tests. Sampling was repeated 50 times, and the mean −log10(*P*) was used for ranking, reducing the influence of differences in segmented-cell number among conditions.

Exact *P* values are reported where shown, and no universal significance threshold was applied except where explicitly indicated. No statistical method was used to predetermine sample size; sample sizes reflected experimental feasibility and the number of analysable cells, lineages, hubs or biological replicates obtained.

### Use of AI-assisted tools

ChatGPT (OpenAI) was used to facilitate the development of custom analysis scripts and to assist with language editing during manuscript preparation. All code, analyses, interpretations and manuscript content were reviewed and verified by the authors.

## Data and material availability

Representative images and corresponding quantitative analyses are provided in the main and Extended Data figures to support evaluation of the reported findings. Previously published imaging data reanalysed in this study are available through the Stanford Digital Repository at https://doi.org/10.25740/ff315yn8920, as described previously^9^.

Raw Hi-C sequencing data are available through GEO under accession GSE330186, with sequencing files deposited in the SRA. Processed files associated with the GEO record include the Juicebox .hic files used for contact-map visualisation and the 25-kb-resolution HiC-Pro .matrix and corresponding _abs.bed files used for ecDNA contact quantification.

Raw RNA-sequencing data are available through GEO under accession GSE348203, with sequencing files deposited in the SRA. Unnormalised gene-level exon and intron count matrices generated using featureCounts, together with the SAF annotation file defining the intronic regions used for quantification, are included as supplementary files associated with the GEO record.

Cell lines and other materials generated or used in this study are available from the corresponding authors upon reasonable request and may require a material transfer agreement.

### Acknowledgements

We thank the members of the Chang and Mischel labs for helpful discussions. We thank Dr. Nathanael S. Gray for sharing the small-molecule compound library and WEE1 degrader. We thank Dr. Anton G. Henssen for sharing the SMS-KAN cell line. We thank Dr. Andrew J. Holland for valuable review and comments, and Dr. Christy Luong for introducing the Hi-C analysis pipeline. We acknowledge the Stanford University Cell Sciences Imaging Core Facility (RRID:SCR_017787) for microscopy support.

## Author contributions

X.Y., P.S.M. and H.Y.C. conceived the project. I.T.W. generated the COLO320DM-FUCCI cells, which X.Y. further engineered to express H2B-emiRFP670. M.M.H. provided the compound library. K.K. trained X.Y. in Hi-C methodology. A.G. performed the metaphase-spread DNA FISH validation of the COLO320DM-TG19 live-cell imaging line. X.Y. purified the RNA, and R.L. prepared the total RNA-sequencing libraries and performed NextSeq sequencing. X.Y. developed hiFISH-96 and all analysis pipelines and performed all other experiments and data analyses reported in the study. X.Y. wrote the original draft, which was reviewed and edited by P.S.M., H.Y.C., I.T.W., A.G. and M.M.H. X.Y., P.S.M. and H.Y.C. acquired funding. P.S.M. and H.Y.C. supervised the project.

---

**Extended Data Fig. 1.**
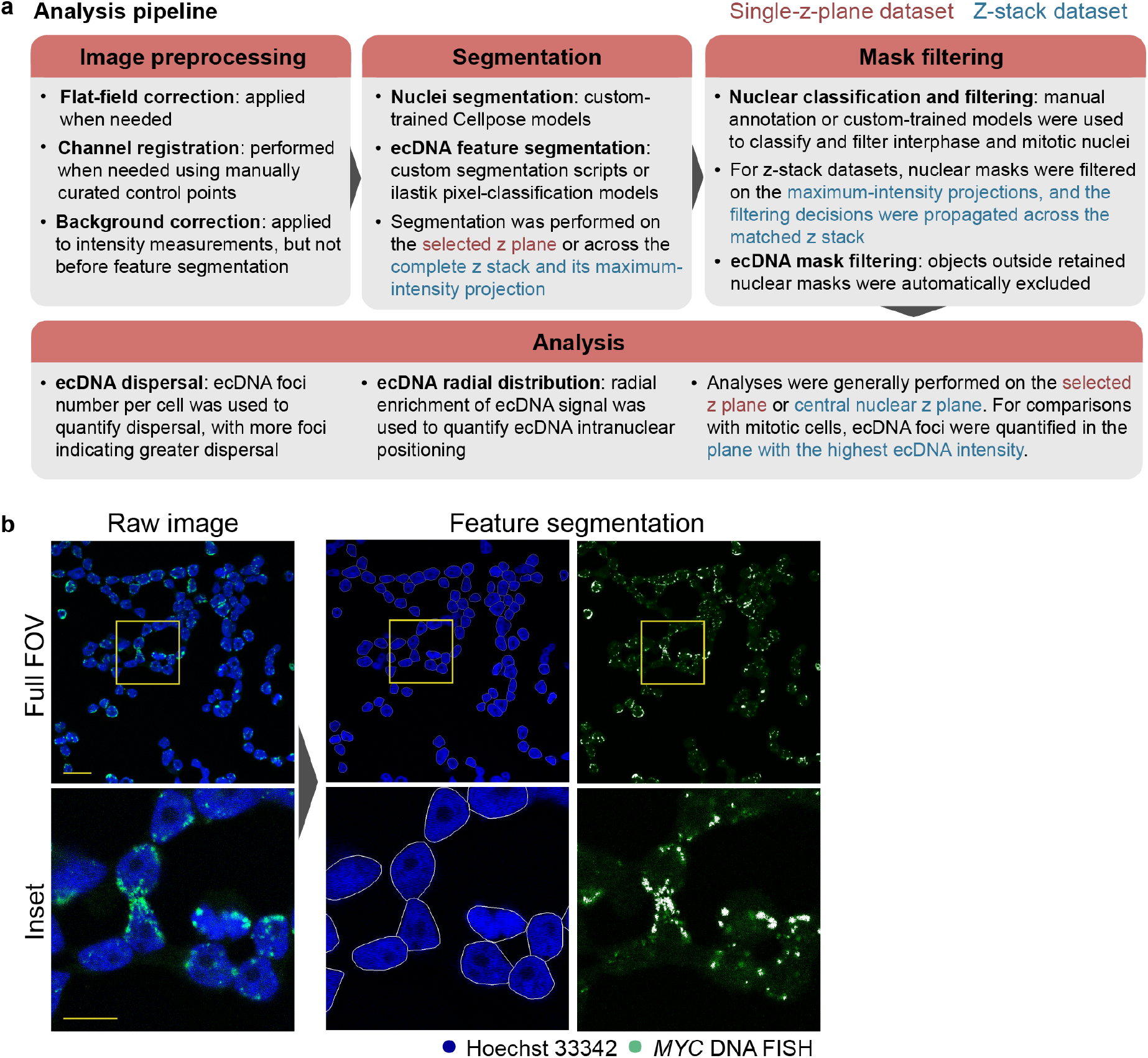
Image-analysis workflow used in the study. **a,** Schematic of the image-analysis pipeline used for preprocessing, segmentation, mask filtering and single-cell quantification of nuclear properties and ecDNA spatial features. Red and blue text indicate procedures specific to single-z-plane and z-stack datasets, respectively. **b,** Representative single-z-plane images from the hiFISH-96 screen shown in Fig. 1. Left, raw images showing Hoechst 33342-stained nuclei (blue) and *MYC* DNA FISH signal marking ecDNA (green). Middle and right, segmentation outputs showing nuclear masks before mask filtering and segmented *MYC* DNA FISH objects before exclusion of objects outside retained nuclear masks. Top, full field of view (FOV); bottom, magnified views of the boxed regions. The workflow enables extraction of object-based ecDNA features from individual nuclei, including ecDNA dispersal, quantified as spatially resolved foci per cell, and radial distribution. Scale bars, 20 μm (top) and 10 μm (bottom).

**Extended Data Fig. 2.**
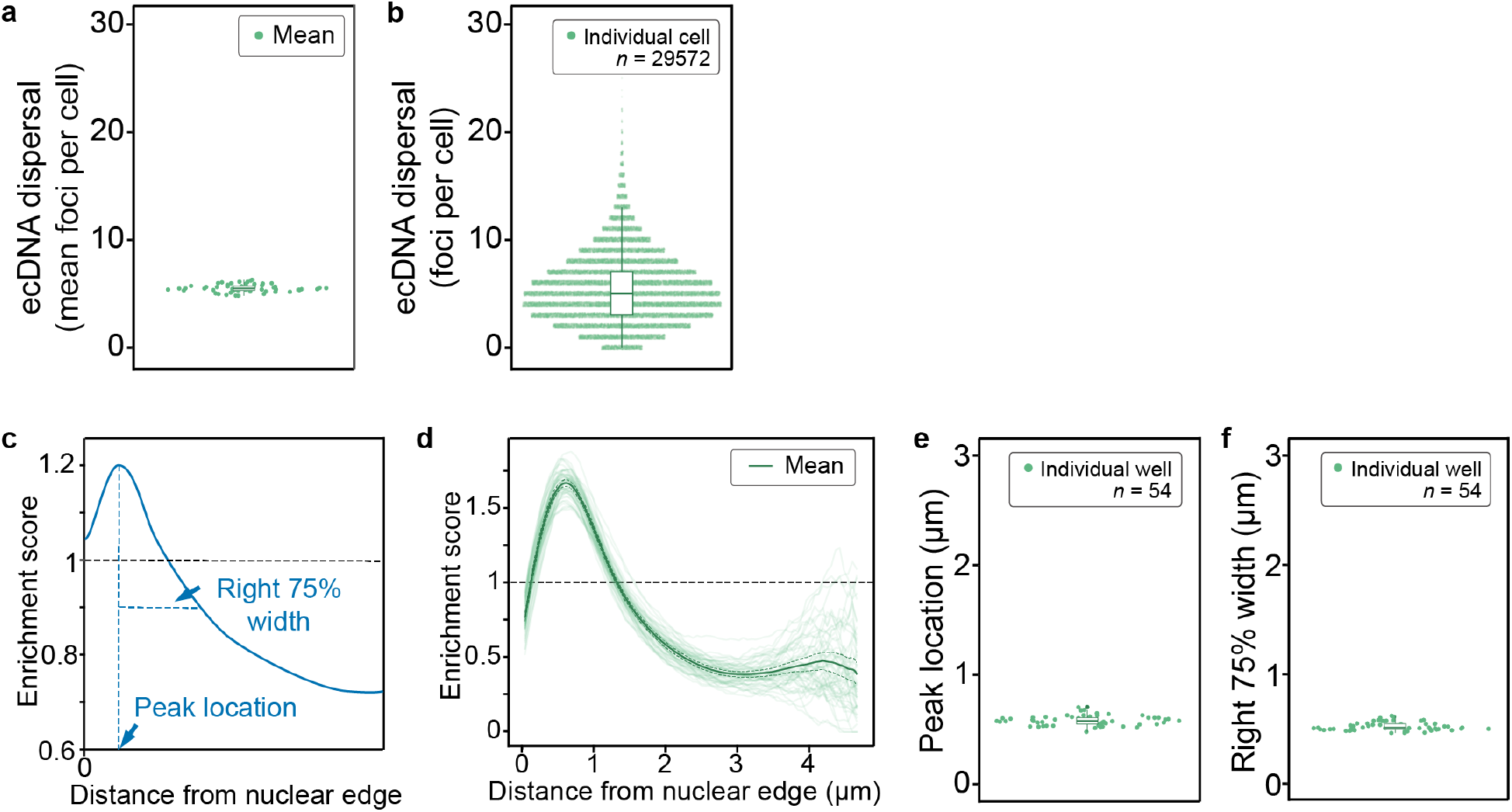
Reproducibility of ecDNA organisation metrics across hiFISH-96 control experiments. DMSO control wells from repeated hiFISH-96 experiments in COLO320DM cells were used to assess the reproducibility of ecDNA organisation metrics. Of 60 DMSO wells, 54 passed all quality-control filters and were retained. **a,** Mean ecDNA dispersal in each retained DMSO well (*n* = 54 wells). **b,** Distribution of ecDNA dispersal across individual cells pooled from the retained wells (*n* = 29,572 cells). **c,** Schematic defining radial enrichment-profile features. Peak location denotes the radial position of maximum enrichment. Right 75% width denotes the distance from the peak to the position on the right-hand side of the profile at which the enrichment score decreases to 75% of its peak value. **d,** Radial enrichment profiles for individual DMSO wells plotted as a function of distance from the nuclear edge, with the mean profile and 95% confidence interval overlaid. The enrichment score was calculated as the fraction of FISH-positive pixels in each radial bin divided by the fraction of total nuclear pixels in that bin; values above or below 1 indicate relative enrichment or depletion, respectively. Horizontal dashed lines in **c,d** denote an enrichment score of 1. **e,f,** Peak location (**e**) and right 75% width (**f**) for each retained DMSO well, as defined in **c**. These analyses demonstrate the reproducibility of multiple ecDNA organisation metrics under baseline conditions and support ecDNA dispersal, quantified as mean foci per cell, as the primary screening metric.

**Extended Data Fig. 3.**
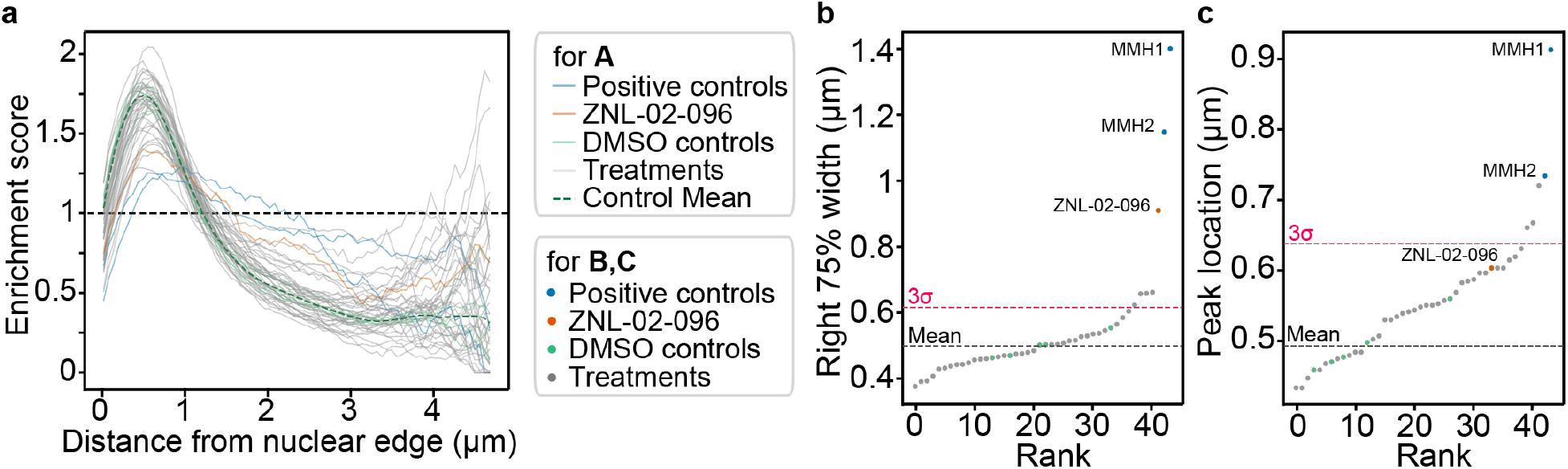
Radial distribution features associated with ecDNA hub dispersal in the hiFISH-96 screen. **a,** Radial enrichment profiles of ecDNA signal in COLO320DM cells across the focused screen, plotted as a function of distance from the nuclear edge. Grey lines indicate individual treatment conditions, light-green lines indicate DMSO controls, blue lines indicate the positive-control BRD4 degraders and the orange line indicates ZNL-02-096. The dashed green line denotes the mean DMSO profile. The enrichment score was calculated as the fraction of FISH-positive pixels in each radial bin divided by the fraction of total nuclear pixels in that bin; values above or below 1 indicate relative enrichment or depletion, respectively. The horizontal black dashed line denotes an enrichment score of 1. **b,c,** Rank-ordered right 75% width (**b**) and peak location (**c**) values for all screened conditions, as defined in Extended Data Fig. 2c. Black dashed lines denote the mean of the DMSO controls included in the screen. Magenta dashed lines denote the DMSO mean plus three standard deviations, with the standard deviation estimated from the repeated DMSO control experiments (*n* = 54 wells). ZNL-02-096 and the positive-control BRD4 degraders shifted the radial distribution of ecDNA relative to DMSO controls, consistent with broader reorganisation of ecDNA positioning accompanying hub dispersal.

**Extended Data Fig. 4.**
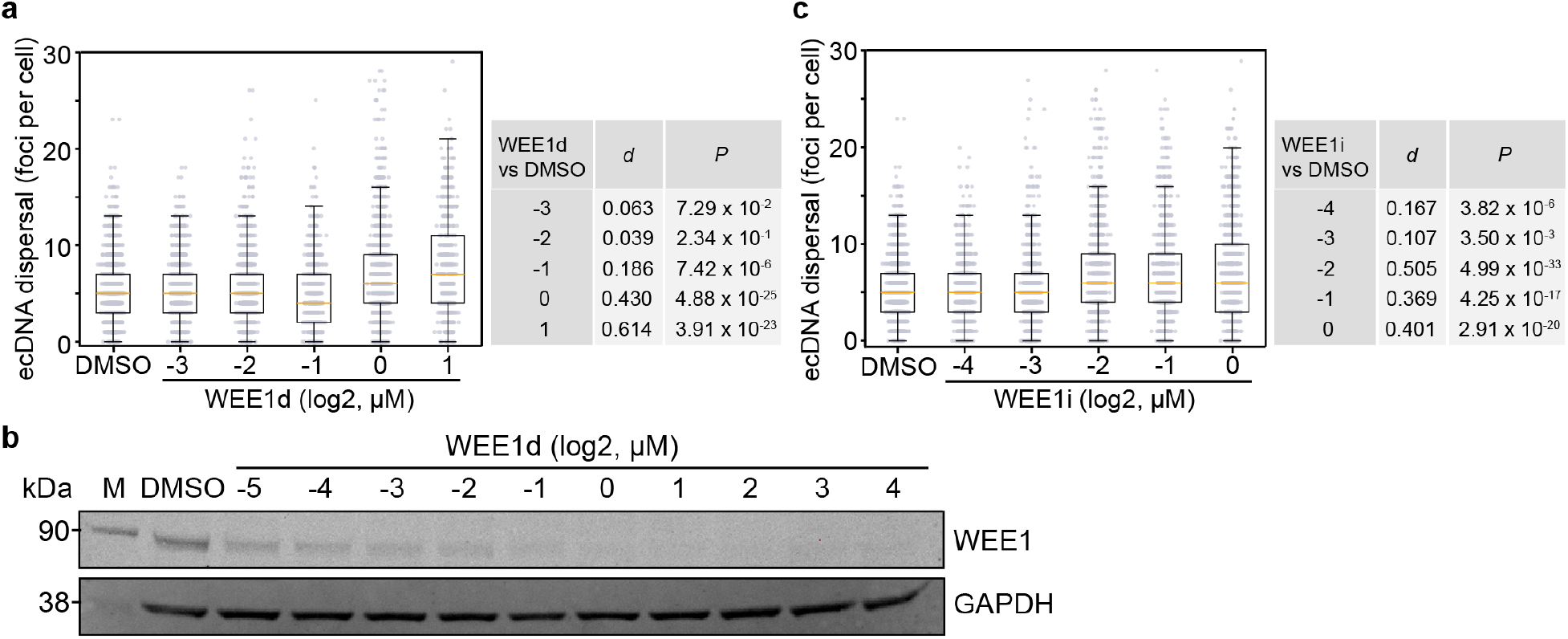
Dose-dependent ecDNA dispersal following WEE1 degradation or inhibition. **a,c,** ecDNA dispersal, quantified as foci per cell, in COLO320DM cells treated for 24 h with increasing concentrations of WEE1d (ZNL-02-096; **a**) or WEE1i (MK-1775; **c**). Both compounds produced dose-dependent increases in ecDNA dispersal. Doses are shown as log2(concentration in μM). Tables show Cohen’s *d* and exact *P* values from two-sided Welch’s *t*-tests comparing each concentration with DMSO. **b,** Immunoblot analysis of WEE1 abundance in COLO320DM cells treated for 24 h with increasing concentrations of WEE1d. WEE1 abundance decreased with increasing WEE1d concentration. GAPDH served as a loading control. In **a,c**, points represent individual cells; box plots show medians and interquartile ranges, and whiskers extend to 1.5× the interquartile range. Sample sizes are provided in Supplementary Table 1.

**Extended Data Fig. 5.**
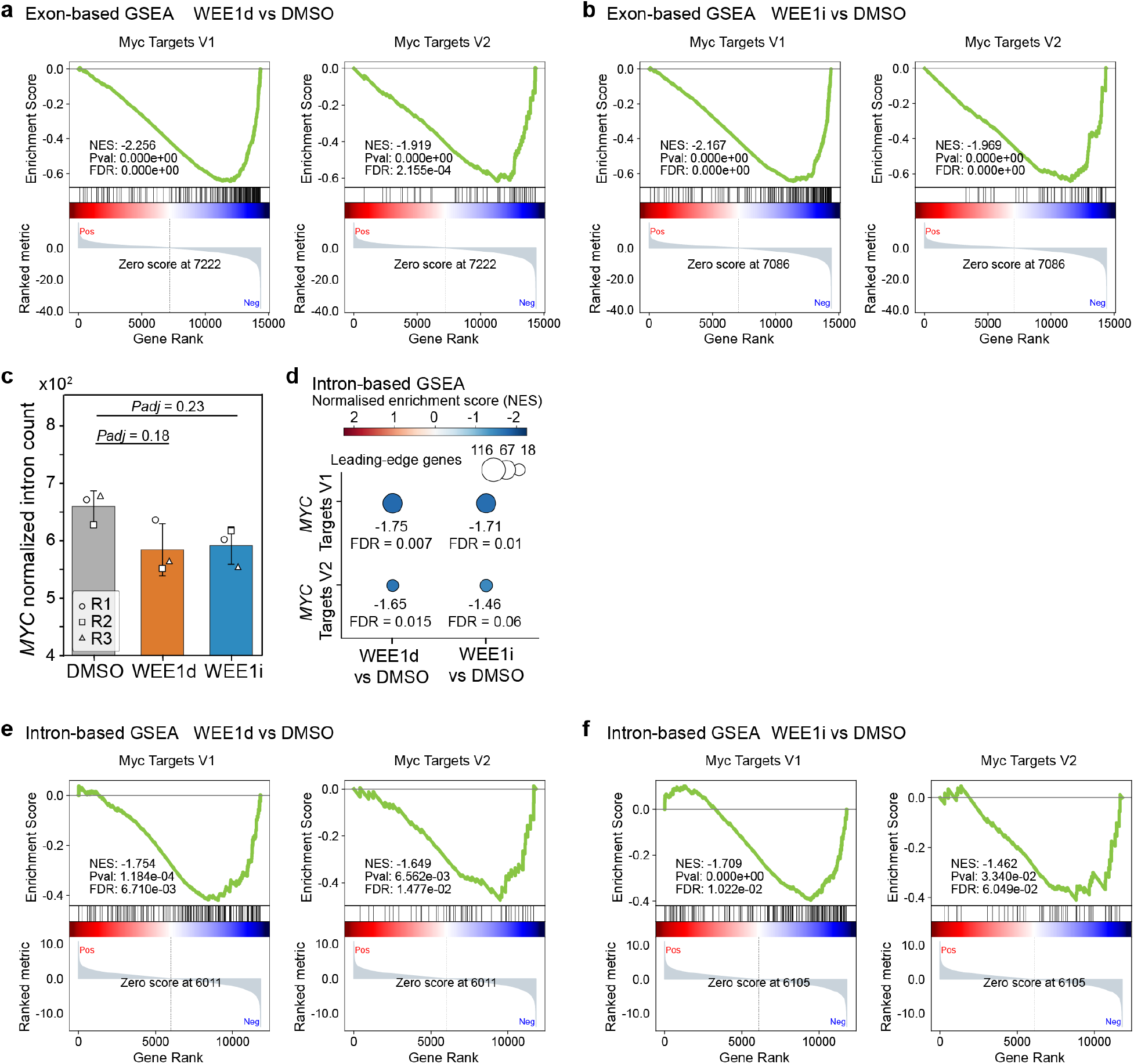
WEE1 perturbation reduces ecDNA-encoded oncogene expression and transcriptional output. Exon-derived reads were used to assess steady-state RNA abundance, whereas intron-derived reads were used as a proxy for nascent transcription. **a,b,** Exon-based GSEA plots for Hallmark MYC Targets V1 and V2 comparing WEE1d with DMSO (**a**) and WEE1i with DMSO (**b**). Green curves show running enrichment scores, black ticks indicate the positions of gene-set members within the ranked lists and the underlying ranking metrics are shown below. Normalised enrichment scores (NES), nominal *P* values and false-discovery rate (FDR) *q* values are indicated. **c,** DESeq2-normalised *MYC* intron counts after 24 h treatment with DMSO, WEE1d or WEE1i (*n* = 3 biological replicates). Bars show means ± s.d.; symbols indicate biological replicates. Adjusted *P* values from DESeq2 Wald tests with Benjamini-Hochberg correction are shown. **d,** Intron-based GSEA of Hallmark MYC Targets V1 and V2 for WEE1d or WEE1i versus DMSO (*n* = 3 biological replicates). Colour indicates NES and bubble size indicates the number of leading-edge genes contributing to enrichment; NES and FDR *q* values are shown. **e,f,** Individual intron-based GSEA plots for Hallmark MYC Targets V1 and V2 comparing WEE1d with DMSO (**e**) and WEE1i with DMSO (**f**), displayed as in **a,b**. Cells were treated with 2 μM WEE1d or 500 nM WEE1i.

**Extended Data Fig. 6.**
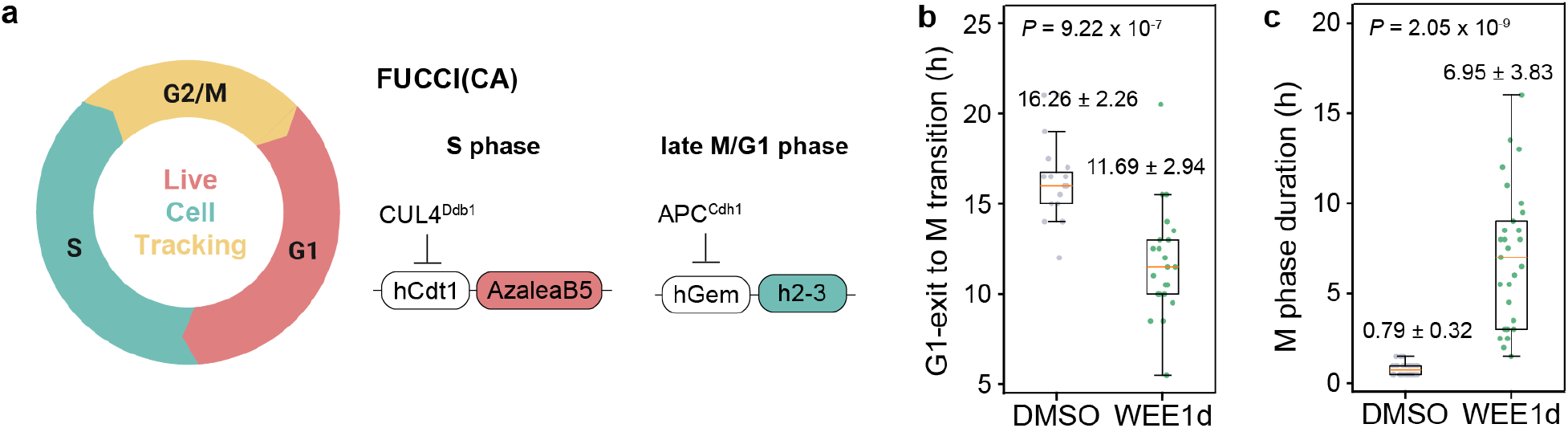
FUCCI(CA) reporter reveals accelerated mitotic entry and delayed mitotic progression in COLO320DM cells. **a–c,** Schematic of the FUCCI(CA) reporter (**a**) and quantification of the interval from G1 exit to mitotic entry (**b**) and M-phase duration (**c**) in DMSO- and WEE1d-treated COLO320DM cells, pooled across *n* = 2 biological replicates. Points represent individual cell lineages; values above the distributions indicate mean ± s.d. Two-sided Welch’s *t*-tests were used in **b,c**, and exact *P* values are shown. Box plots show medians and interquartile ranges; whiskers extend to 1.5× the interquartile range. Lineage numbers are provided in Supplementary Table 1.

**Extended Data Fig. 7.**
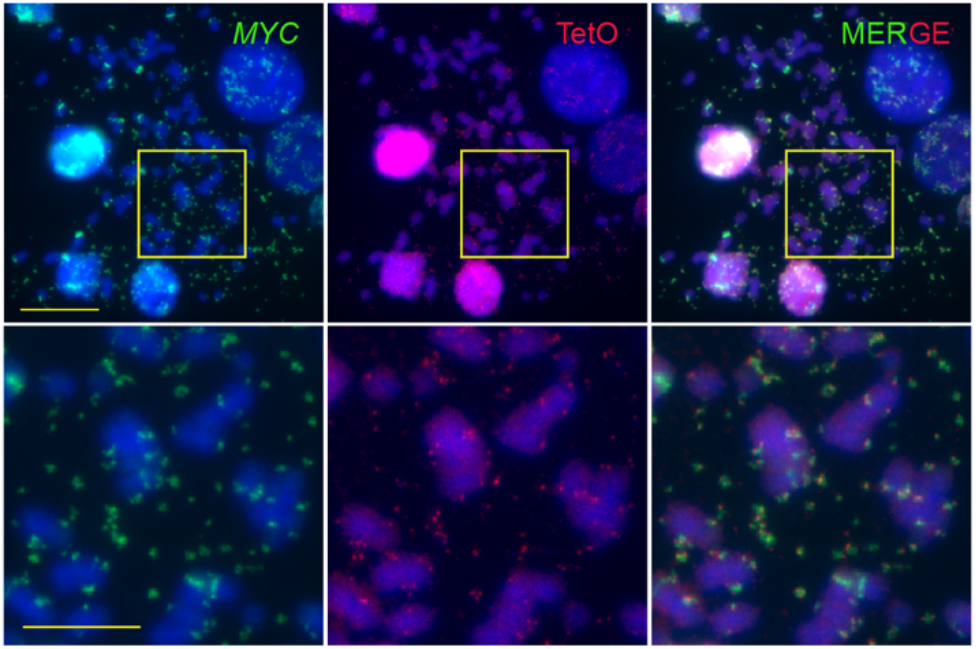
Metaphase-spread validation of the *MYC* ecDNA live-cell imaging line. Representative metaphase-spread DNA FISH images of the engineered COLO320DM-TG19 line used for live-cell tracking of ecDNA dynamics. *MYC* DNA FISH signal is shown in green, TetO DNA FISH signal in red and Hoechst 33342-stained DNA in blue. Merged images show colocalisation of the TetO array with *MYC*-containing ecDNA, supporting the locus-labelling strategy used for live-cell imaging. Top, full field of view; bottom, magnified views of the boxed regions. Scale bars, 20 μm (top) and 10 μm (bottom).

**Extended Data Fig. 8.**
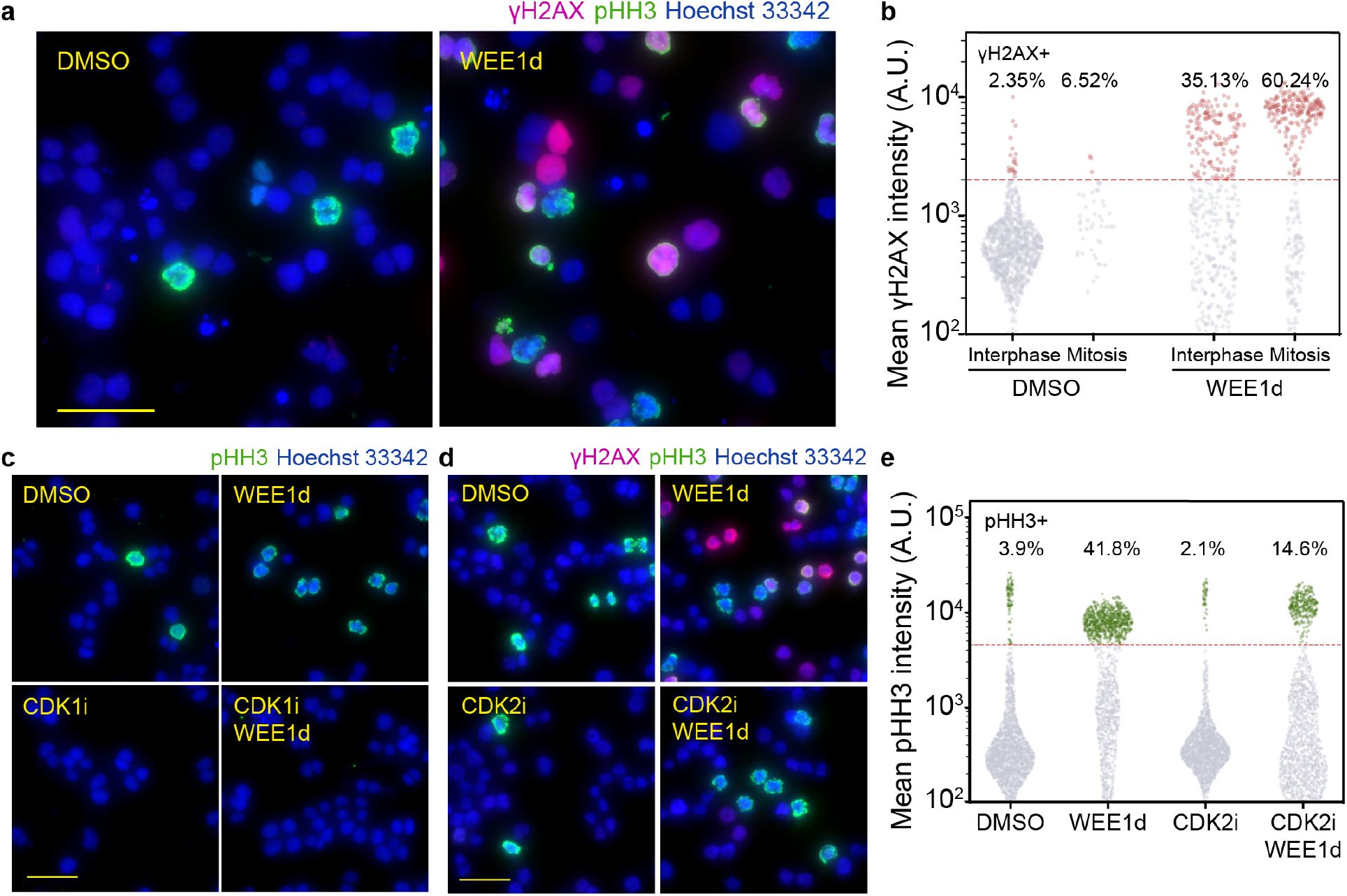
DNA-damage signalling and mitotic entry following WEE1 degradation. **a,b,** Representative immunofluorescence images (**a**) and mean nuclear γH2AX intensity (**b**) in interphase and mitotic COLO320DM cells treated for 6 h with DMSO or WEE1d. γH2AX is magenta, phospho-histone H3 (pHH3) is green and Hoechst 33342-stained nuclei are blue. Cells were classified as mitotic or interphase according to pHH3 positivity. In **b**, points represent individual nuclei, the red dashed line denotes the threshold for γH2AX positivity and percentages indicate the fraction of nuclei above this threshold. The increase in γH2AX among WEE1d-treated interphase cells supports the induction of DNA-damage signalling before mitotic entry. **c,** Representative pHH3 immunofluorescence images of cells treated for 6 h with DMSO, WEE1d, CDK1i (RO-3306) or CDK1i plus WEE1d. pHH3 is green and nuclei are blue. Quantification is shown in Fig. 4e. **d,e,** Representative γH2AX and pHH3 immunofluorescence images (**d**) and mean nuclear pHH3 intensity (**e**) after 6 h of treatment with DMSO, WEE1d, CDK2i (BLU-222) or CDK2i plus WEE1d. γH2AX is magenta, pHH3 is green and nuclei are blue. In **e**, points represent individual nuclei, the red dashed line denotes the threshold for pHH3 positivity and percentages indicate the fraction of nuclei above this threshold. γH2AX quantification corresponding to **d** is shown in Fig. 4h. CDK2 inhibition reduced WEE1d-induced γH2AX signalling while retaining a pHH3-positive population, indicating that DNA-damage signalling was suppressed without completely blocking mitotic entry. Within each experiment, positivity thresholds were applied consistently across treatment groups. Data are from *n* = 3 biological replicates. Concentrations: 2 μM WEE1d, 10 μM CDK1i and 125 nM CDK2i. Cell numbers are provided in Supplementary Table 1. Scale bars, 40 μm.

**Extended Data Fig. 9.**
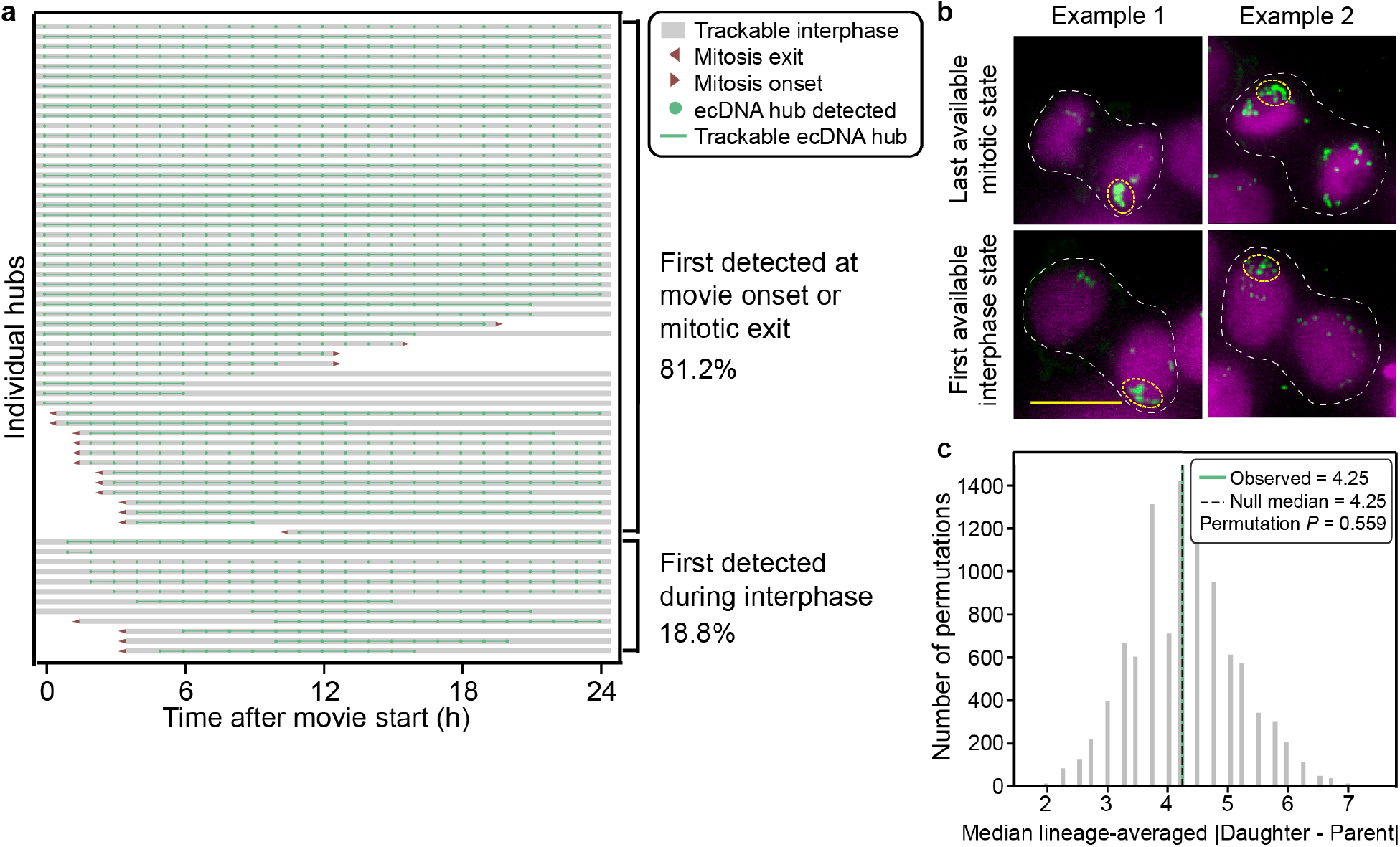
Interphase ecDNA hubs persist, whereas mitosis enables remodelling. **a,** Tracking summary for ecDNA hubs in DMSO-treated COLO320DM cells (*n* = 4 biological replicates). Hubs larger than 2 μm² were manually tracked in 24-h movies acquired hourly; only interphases longer than 12 h were retained. Rows represent individual hubs; green points indicate detection, green lines indicate trackable duration, grey bars denote trackable interphase, and arrowheads mark mitotic onset or exit. Hubs were grouped according to whether they were first detected at movie onset or mitotic exit (81.2%) or later during interphase (18.8%). Similar proportions were observed when the analysis was restricted to interphases tracked from a clearly observed mitotic exit (Fig. 5c). **b,** Examples of paired last mitotic and first daughter-interphase states from matched lineages. Dashed white outlines mark tracked lineages, and yellow circles highlight representative ecDNA hubs. **c,** Permutation analysis of parent-daughter similarity. The observed median lineage-averaged absolute parent-daughter difference was compared with a null distribution generated by random reassignment of daughter identities; the observed value, null median and permutation *P* value are shown. Images in **b** are SVCC-rendered maximum-intensity projections; ecDNA is green and chromosomal DNA is magenta. Sample sizes are provided in Supplementary Table 1. Scale bars, 10 μm.

**Extended Data Fig. 10.**
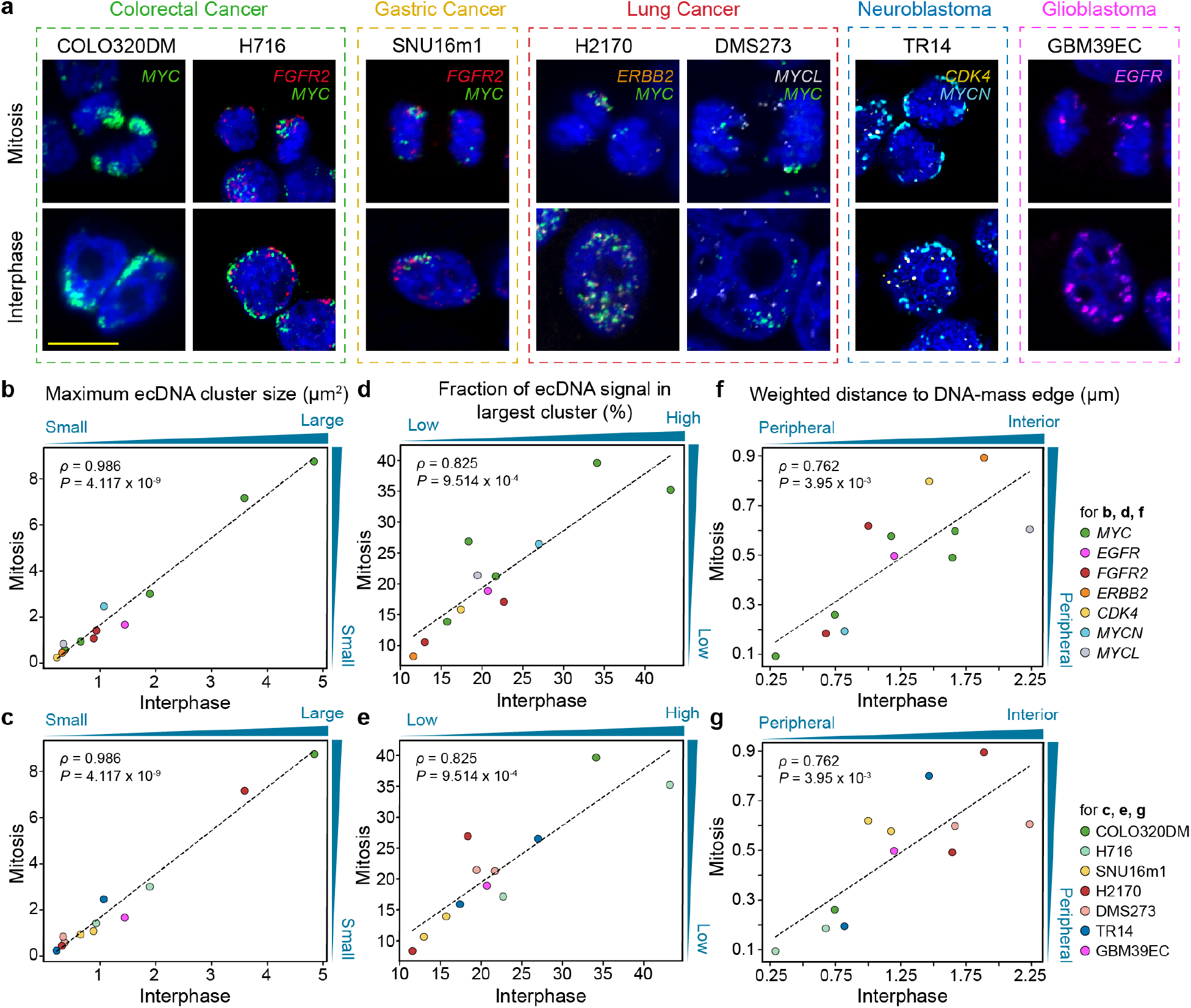
Mitotic and interphase ecDNA organisation correspond across cancer models. **a,** Representative mitotic and interphase DNA FISH images from seven ecDNA-positive cell lines spanning colorectal, gastric and lung cancers, neuroblastoma and glioblastoma. FISH signals are coloured as indicated; Hoechst 33342 or DAPI is blue. Images of H716 and TR14 cells were obtained from previously generated DNA FISH datasets^9^ and reanalysed here. **b–e,** Comparison of interphase and mitotic ecDNA organisation using maximum ecDNA cluster size (**b,c**) and the fraction of ecDNA signal within the largest cluster (**d,e**). **f,g,** Comparison of the intensity-weighted distance of ecDNA signal from the DNA-mass edge in interphase and mitotic populations. Larger values indicate more internally positioned ecDNA relative to the DNA-mass edge. In **b–g**, points represent cell line-oncogene pairs and are coloured by ecDNA-associated oncogene (**b,d,f**) or cell line (**c,e,g**); dashed lines denote *y* = *x*. Associations were assessed by two-sided Spearman correlation; correlation coefficients and exact *P* values are shown. Sample sizes are provided in Supplementary Table 2. Scale bars, 10 μm.

## Supplementary Information

This PDF contains Supplementary Tables 1–9 describing imaging sample sizes, cross-model analyses, culture conditions, experimental workflows, probes, imaging settings, antibodies and reagents.

## Supplementary Data 1

Compound-level results from the hiFISH-96 imaging screen.

**Supplementary Table 1.**
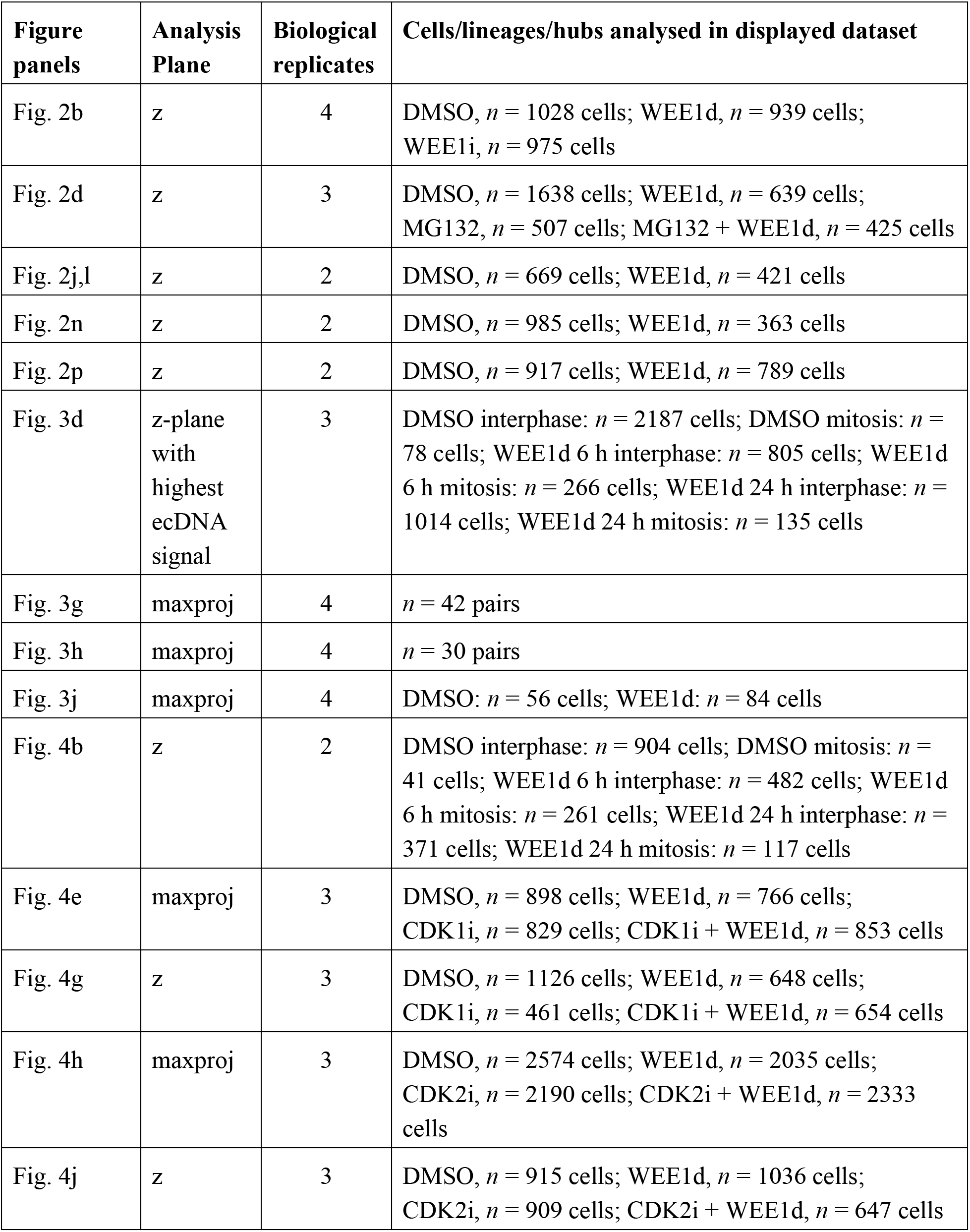
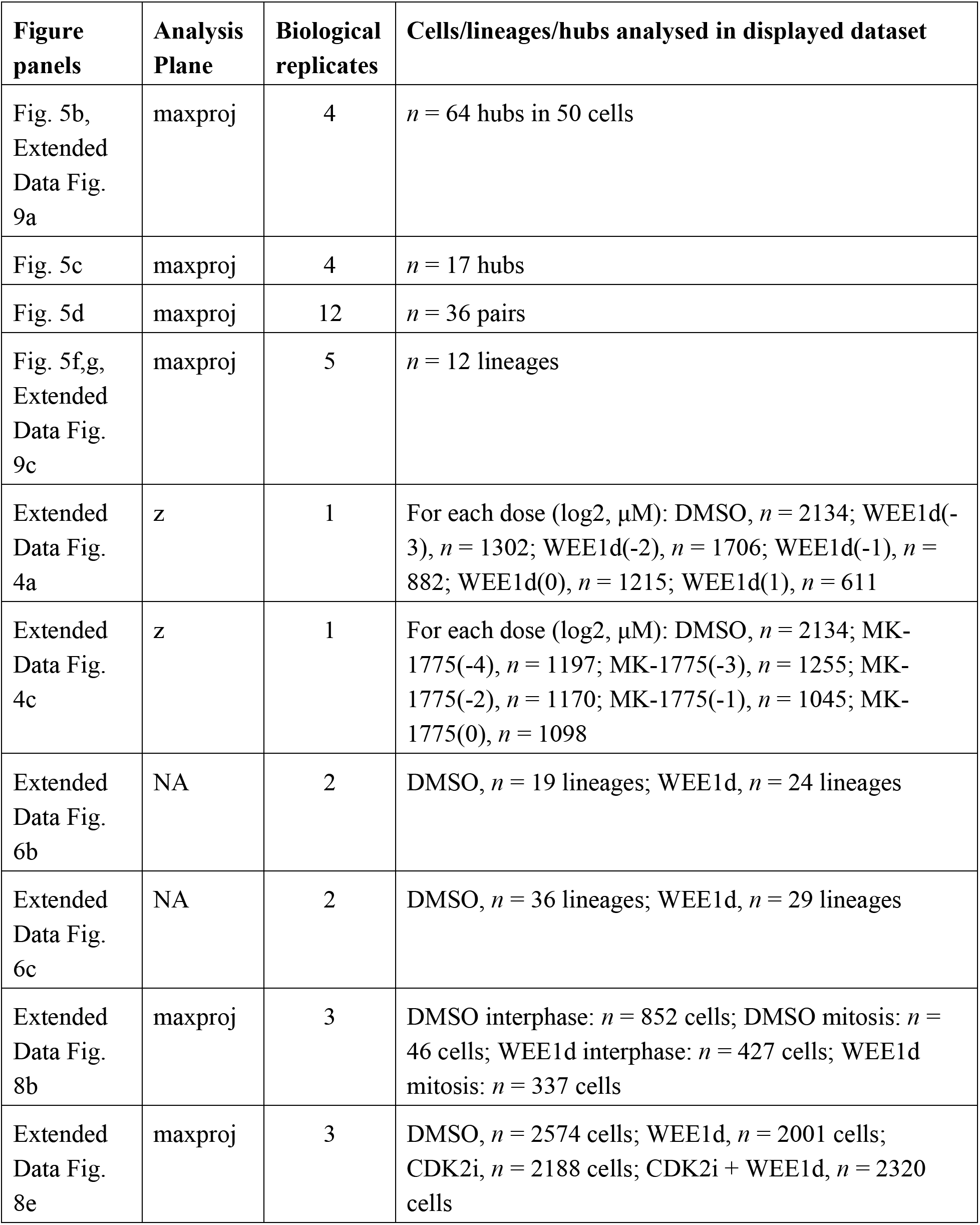
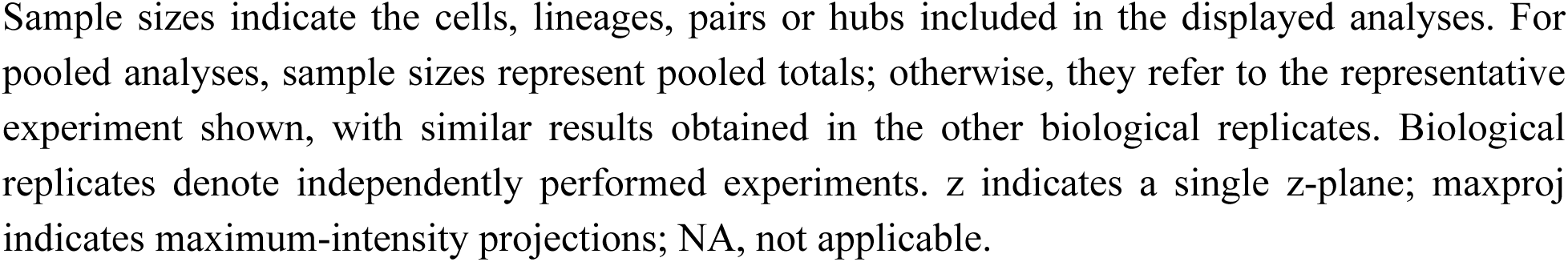
Biological replicates and sample sizes for imaging analyses. Sample sizes indicate the cells, lineages, pairs or hubs included in the displayed analyses. For pooled analyses, sample sizes represent pooled totals; otherwise, they refer to the representative experiment shown, with similar results obtained in the other biological replicates. Biological replicates denote independently performed experiments. z indicates a single z-plane; maxproj indicates maximum-intensity projections; NA, not applicable.

**Supplementary Table 2.**
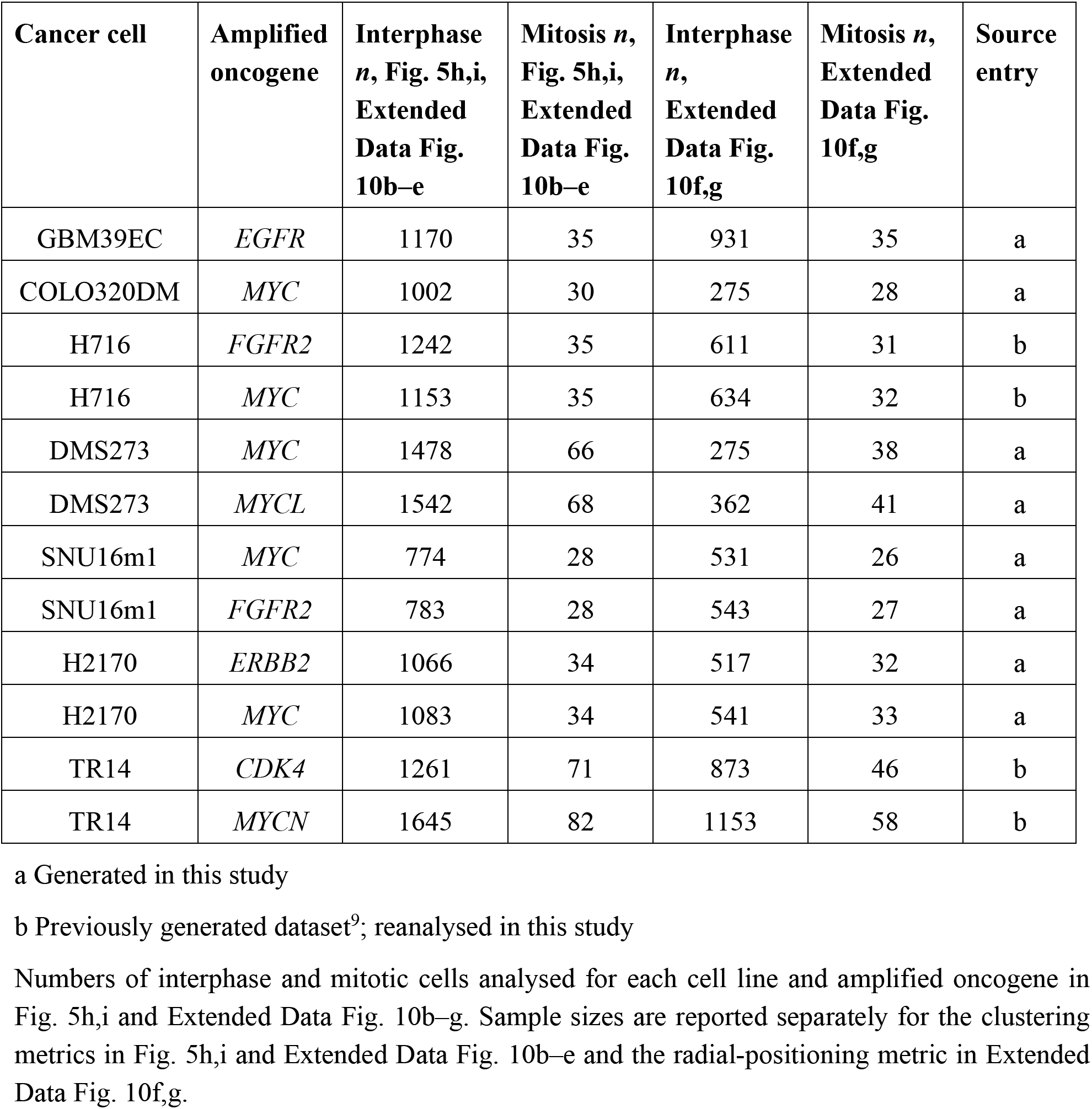
Sample sizes for cross-cell-line analyses of mitotic and interphase ecDNA organisation. Numbers of interphase and mitotic cells analysed for each cell line and amplified oncogene in Fig. 5h,i and Extended Data Fig. 10b–g. Sample sizes are reported separately for the clustering metrics in Fig. 5h,i and Extended Data Fig. 10b–e and the radial-positioning metric in Extended Data Fig. 10f,g.

**Supplementary Table 3.** Cell lines and culture conditions.

| **Cell line** | **Source, identifier and derivation** | **Culture medium and supplements** |
| --- | --- | --- |
| COLO320DM | ATCC, CCL-220,<br>RRID:CVCL_0219 | maintained in DMEM supplemented with 10% FBS and 1% PSG |
| H2170 | ATCC, CRL-5928,<br>RRID:CVCL_1535 | Maintained in ATCC-formulated RPMI 1640 medium supplemented with 10% FBS and 1% PSG |
| H716 | ATCC, CCL-251,<br>RRID:CVCL_1581 | Maintained in ATCC-formulated RPMI 1640 medium supplemented with 10% FBS and 1% PSG |
| DMS273 | Sigma-Aldrich, 95062830-1VL,<br>RRID:CVCL_1176 | Maintained in ATCC-formulated RPMI 1640 medium supplemented with 10% FBS and 1% PSG |
| SMS-KAN | gift from the laboratory of Anton G. Henssen, RRID:CVCL_7131 | Maintained in ATCC-formulated RPMI 1640 medium supplemented with 10% FBS and 1% PSG |
| TR14 | provided by J. J. Molenaar at the Princess Máxima Center for Pediatric Oncology as described previously<sup>9</sup>. | Maintained in RPMI 1640 medium supplemented with 20% FBS and 1% PS. |
| SNU16m1 | Monoclonal SNU16m1 line was derived from parental SNU16 cells by single-cell cloning following lentiviral transduction to stably express dCas9-KRAB, as described previously<sup>6</sup>. | Maintained in DMEM/F-12 1:1, supplemented with 10% FBS and 1% PSG |
| GBM39EC | human patient-derived glioblastoma neurosphere line established as described previously<sup>7,33</sup>. | Maintained in DMEM/F12 1:1 supplemented with B-27 supplement, 1% PS, 1% GlutaMAX, 20 ng ml<sup>-1</sup> hEGF, 20 ng ml<sup>-1</sup> hFGF and 5 µg ml<sup>-1</sup> heparin |

| Cell line | Source, identifier and derivation | Culture medium and supplements |
| --- | --- | --- |
| COLO320DM-TG19 | COLO320DM-TG19, which contains a TetO array adjacent to *MYC*-containing ecDNA and stably expresses TetR-mNeonGreen and H2B-emiRFP670, was generated from COLO320DM cells as described previously<sup>9</sup>. | Same as COLO320DM |
| COLO320DM-FUCCI H2B-emiRFP670 | Generated and described in this study | Same as COLO320DM |

FBS: fetal bovine serum; PS: penicillin-streptomycin; PSG: penicillin-streptomycin-glutamine; hEGF: human epidermal growth factor; hFGF: human fibroblast growth factor

**Supplementary Table 4.** Cell coating reagents and methods used in this study.

| **Coating Reagents** | **Coating Method** | **Used for** |
| --- | --- | --- |
| Poly-D-lysine | 10 $\mu\text{g ml}^{-1}$ poly-D-lysine (Sigma-Aldrich, A-003-E) was applied for 1 h at 37 °C, coverslips or glass-bottom imaging plates were washed with PBS and allowed to dry before cell seeding | COLO320DM, SNU16m1, H2170, DMS273, SMS-KAN, COLO320DM-TG19 |
| Fibronectin | 10 $\mu\text{g ml}^{-1}$ fibronectin (Sigma-Aldrich, FC010-5MG) was applied for 1 h at 37 °C, coverslips or glass-bottom imaging plates were washed with PBS and allowed to dry before cell seeding | COLO320DM |
| Laminin | 10 $\mu\text{g ml}^{-1}$ laminin (Sigma-Aldrich, L2020) was applied for 1 h at 37 °C, coverslips or glass-bottom imaging plates were washed with PBS and allowed to dry before cell seeding | GBM39EC |

**Supplementary Table 5.** DNA FISH methods used in this study.

| **Input samples** | **Methods** | **Used for** |
| --- | --- | --- |
| Cells seeded on coated coverslip | Heat-denaturation method | General coverslip-based DNA FISH |
|  | Acid-denaturation method | DNA FISH following immunofluorescence, or used separately |
| Cells seeded in coated 96-well glass-bottom imaging plates | hiFISH-96 | General high-throughput DNA FISH |
| Metaphase spread sample | Metaphase DNA FISH | DNA FISH following metaphase spread |

**Supplementary Table 6.**
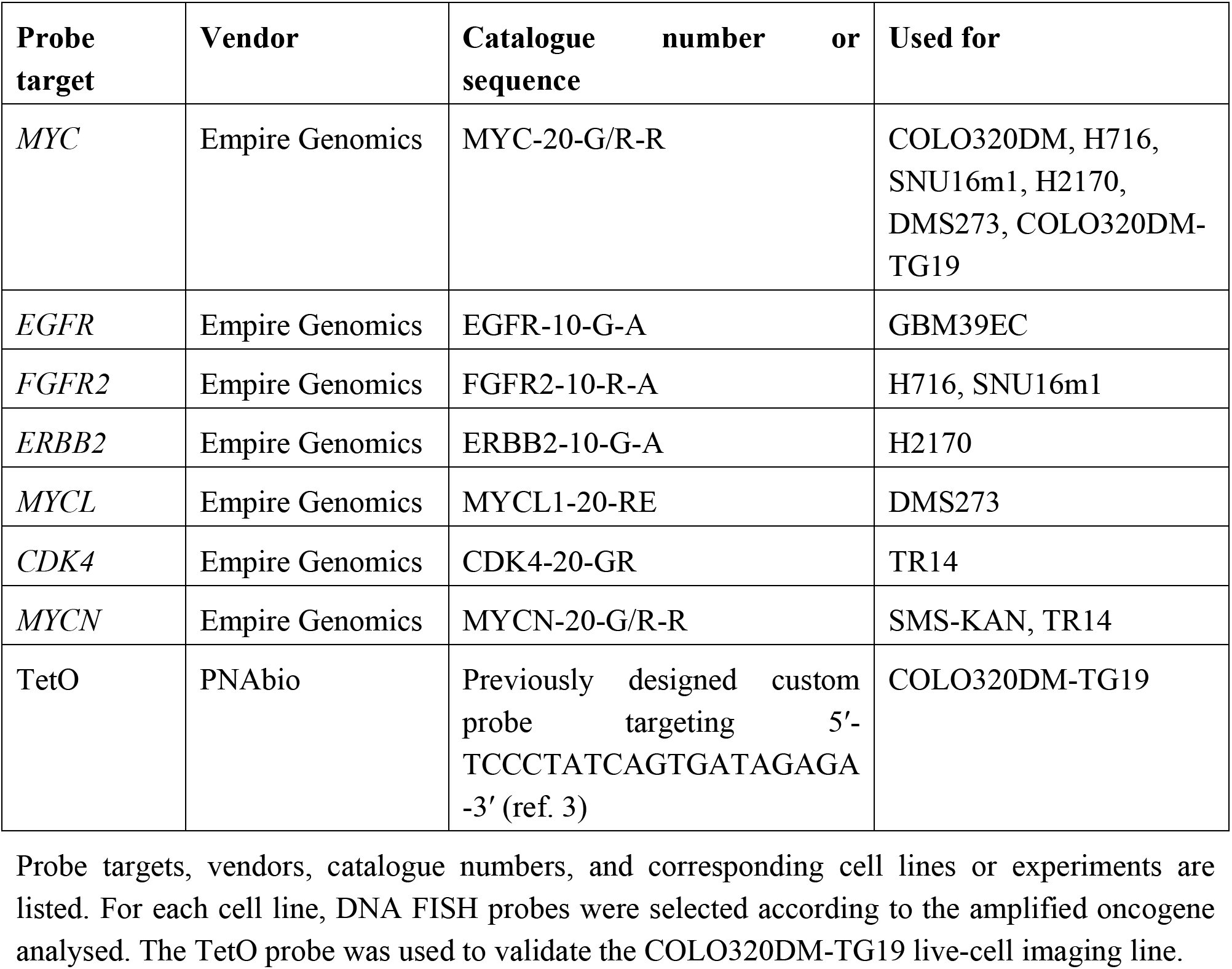
DNA FISH probes used in this study. Probe targets, vendors, catalogue numbers, and corresponding cell lines or experiments are listed. For each cell line, DNA FISH probes were selected according to the amplified oncogene analysed. The TetO probe was used to validate the COLO320DM-TG19 live-cell imaging line.

**Supplementary Table 7.** Antibodies used in the study.

| **Antibody** | **Source** | **Identifier** | **Dilution used** |
| --- | --- | --- | --- |
| rabbit anti-pHH3 antibody | Abcam, ab5176 | RRID:AB_304763 | 1:300 |
| mouse anti-γH2AX antibody | Sigma-Aldrich, 05-636 | RRID:AB_309864 | 1:300 |
| Alexa Fluor 488 goat anti-rabbit IgG | Thermo Fisher Scientific, A11008 | RRID:AB_143165 | 1:500 |
| Alexa Fluor 647 goat anti-rabbit IgG | Thermo Fisher Scientific, A21245 | RRID:AB_2535813 | 1:500 |
| Alexa Fluor 555 goat anti-mouse IgG | Thermo Fisher Scientific, A21422 | RRID:AB_141822 | 1:500 |
| anti-WEE1 antibody | Cell Signaling, 4936S | RRID:AB_2288509 | 1:100 |
| anti-GAPDH antibody | Abcam, ab8245 | RRID:AB_2107448 | 1:1000 |
| IRDye® 800CW Goat anti-Rabbit IgG Secondary Antibody | LICORbio, 926-32211 | RRID:AB_621843 | 1:15,000 |
| IRDye® 680RD Goat Anti-Mouse IgG Secondary Antibody | LICORbio, 926-68070 | RRID:AB_10956588 | 1:15,000 |

**Supplementary Table 8.**
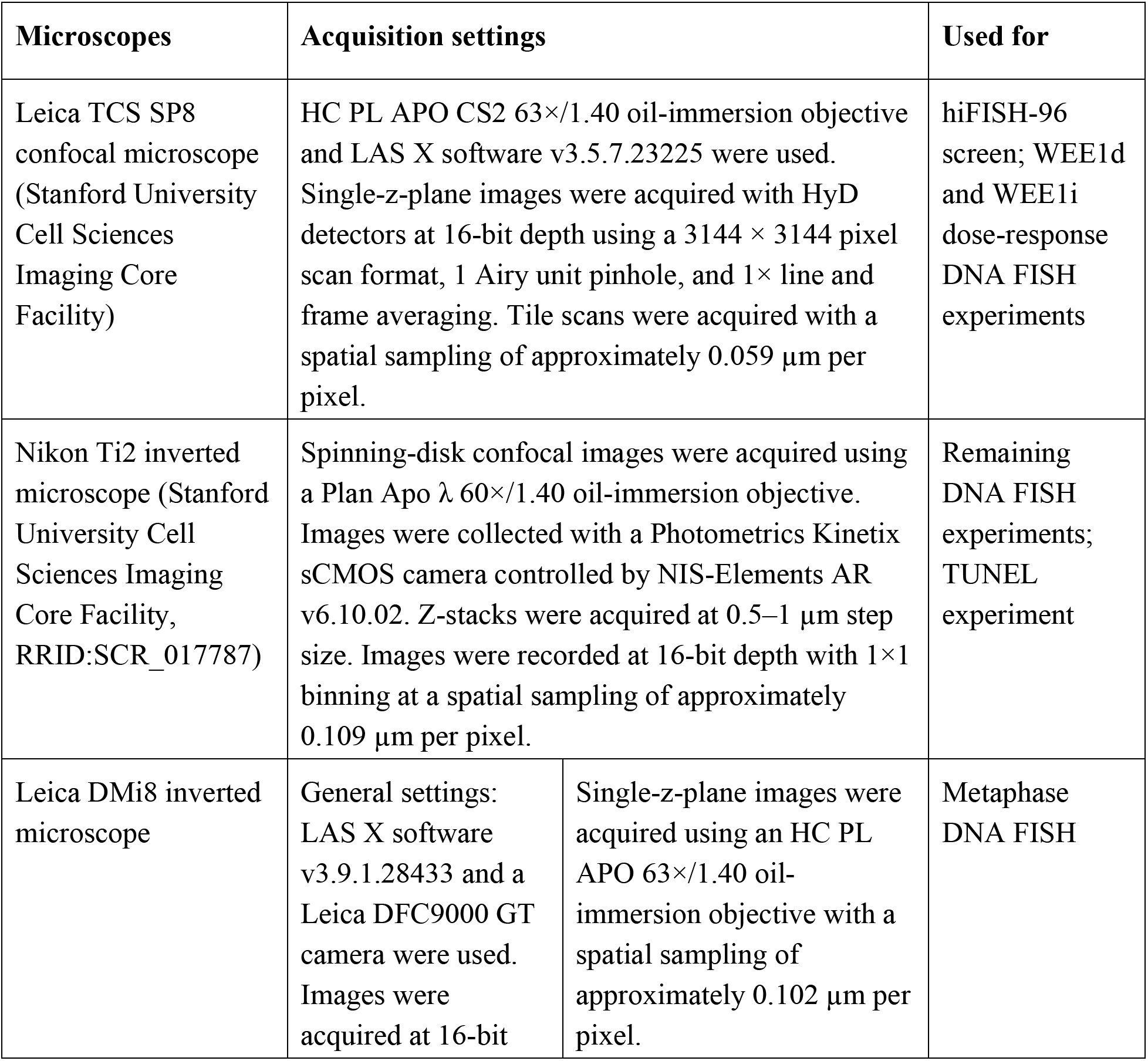
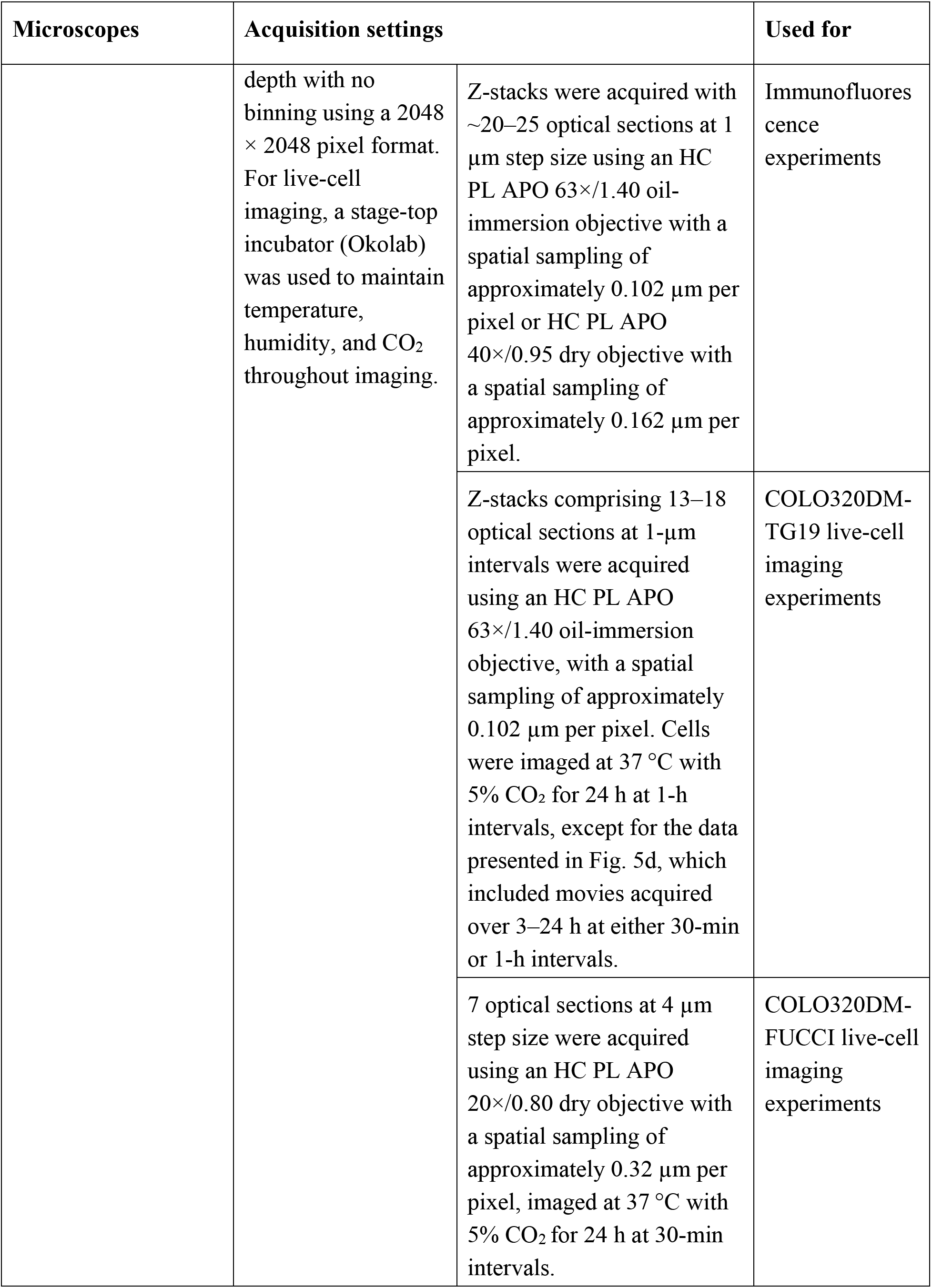
Imaging acquisition conditions used in the study.

| **Microscopes** | **Acquisition settings** |  | **Used for** |
| --- | --- | --- | --- |
| Leica TCS SP8 confocal microscope (Stanford University Cell Sciences Imaging Core Facility) | HC PL APO CS2 63×/1.40 oil-immersion objective and LAS X software v3.5.7.23225 were used. Single-z-plane images were acquired with HyD detectors at 16-bit depth using a 3144 × 3144 pixel scan format, 1 Airy unit pinhole, and 1× line and frame averaging. Tile scans were acquired with a spatial sampling of approximately 0.059 μm per pixel. |  | hiFISH-96 screen; WEE1d and WEE1i dose-response DNA FISH experiments |
| Nikon Ti2 inverted microscope (Stanford University Cell Sciences Imaging Core Facility, RRID:SCR_017787) | Spinning-disk confocal images were acquired using a Plan Apo λ 60×/1.40 oil-immersion objective. Images were collected with a Photometrics Kinetix sCMOS camera controlled by NIS-Elements AR v6.10.02. Z-stacks were acquired at 0.5–1 μm step size. Images were recorded at 16-bit depth with 1×1 binning at a spatial sampling of approximately 0.109 μm per pixel. |  | Remaining DNA FISH experiments; TUNEL experiment |
| Leica DMI8 inverted microscope | General settings: LAS X software v3.9.1.28433 and a Leica DFC9000 GT camera were used. Images were acquired at 16-bit | Single-z-plane images were acquired using an HC PL APO 63×/1.40 oil-immersion objective with a spatial sampling of approximately 0.102 μm per pixel. | Metaphase DNA FISH |

| Microscopes | Acquisition settings |  | Used for |
| --- | --- | --- | --- |
|  | depth with no binning using a 2048 × 2048 pixel format. For live-cell imaging, a stage-top incubator (Okolab) was used to maintain temperature, humidity, and CO<sub>2</sub> throughout imaging. | Z-stacks were acquired with ~20–25 optical sections at 1 μm step size using an HC PL APO 63×/1.40 oil-immersion objective with a spatial sampling of approximately 0.102 μm per pixel or HC PL APO 40×/0.95 dry objective with a spatial sampling of approximately 0.162 μm per pixel. | Immunofluorescence experiments |
|  |  | Z-stacks comprising 13–18 optical sections at 1-μm intervals were acquired using an HC PL APO 63×/1.40 oil-immersion objective, with a spatial sampling of approximately 0.102 μm per pixel. Cells were imaged at 37 °C with 5% CO<sub>2</sub> for 24 h at 1-h intervals, except for the data presented in Fig. 5d, which included movies acquired over 3–24 h at either 30-min or 1-h intervals. | COLO320DM-TG19 live-cell imaging experiments |
|  |  | 7 optical sections at 4 μm step size were acquired using an HC PL APO 20×/0.80 dry objective with a spatial sampling of approximately 0.32 μm per pixel, imaged at 37 °C with 5% CO<sub>2</sub> for 24 h at 30-min intervals. | COLO320DM-FUCCI live-cell imaging experiments |

**Supplementary Table 9.** Additional reagents and materials used in the study. Reagents and materials used for DNA FISH, hiFISH-96, metaphase spread, immunofluorescence, live-cell imaging, Hi-C, immunoblotting, TUNEL analysis and RNA sequencing (RNA-seq) are listed with suppliers and catalogue numbers.

| **Reagent/kit/material** | **Vendor** | **Catalogue number** |
| --- | --- | --- |
| Dulbecco's modified Eagle's medium/nutrient mixture F-12 (DMEM/F-12 1:1) | Gibco | 11320-033 |
| Fetal bovine serum (FBS) | Hyclone | SH30396.03 |
| Penicillin-streptomycin-glutamine (PSG) | Thermo Fisher Scientific | 10378016 |
| Penicillin-streptomycin (PS) | Thermo Fisher Scientific | 15140-122 |
| DMEM | Corning | 10-013-CV |
| B-27 supplement | Gibco | 17504044 |
| GlutaMAX | Gibco | 35050061 |
| human epidermal growth factor (hEGF) | Sigma-Aldrich | E9644 |
| human fibroblast growth factor (hFGF) | Sigma-Aldrich | F0291 |
| heparin | Sigma-Aldrich | H3149-500KU |
| TrypLE Express | Gibco | 12605010 |
| ATCC-formulated RPMI 1640 medium | Gibco | A1049101 |
| RPMI 1640 medium | Gibco | 22400089 |
| MycoAlert PLUS Mycoplasma Detection Kit | Lonza | LT07-710 |
| tFucci(CA)5 plasmid | Addgene | 153521 |
| Super PiggyBac transposase expression vector | System Biosciences | PB210PA-1 |
| Opti-MEM | Gibco | 31985062 |
| X-tremeGENE™ HP DNA Transfection Reagent | Roche | 6366236001 |
| TransIT-LT1 Transfection Reagent | Mirus Bio | MIR 2300 |
| polybrene | EMD Millipore | TR-1003-G |
| HEPES | Gibco | 15630080 |
| MK-1775 (WEE1i) | Selleck Chemicals | S1525 |
| BLU-222 (CDK2i) | MedChemExpress | HY-160701 |
| MG132 | Cayman Chemical | 10012628 |
| RO-3306 (CDK1i) | Cayman Chemical | 15149 |
| Paraformaldehyde (PFA) | Thermo Fisher Scientific/Electron Microscopy Sciences | 50-980-487/15710-S |
| Triton X-100 | Sigma-Aldrich | T8787 |
| Hydrochloric acid (HCl) | Thermo Fisher Scientific | A144-500 |
| 20× SSC | Ambion | AM9770 |
| Tween-20 | Sigma-Aldrich | 11332465001 |
| Hoechst 33342 | Invitrogen | H3570 |
| DAPI | Sigma-Aldrich | D9542 |
| ProLong Gold Antifade Mountant | Invitrogen | P36935 |
| ProLong Diamond Antifade Mountant | Invitrogen | P36965 |
| Rubber cement | MP Biomedicals | MP1FIXO0125 |
| KaryoMAX Colcemid | Gibco | 15212012 |
| KCl | Ambion | AM9640G |
| Methanol | Thermo Fisher Scientific | A412-1 |
| Acetic acid | Thermo Fisher Scientific | BP1185-500 |
| BSA | Sigma-Aldrich | A2153-50G |
| Click-iT TUNEL Alexa Fluor Imaging Assay for Microscopy and HCS | Thermo Fisher Scientific | C10245 |
| ProLong Live Antifade Reagent | Thermo Fisher Scientific | P36975 |
| Formaldehyde | Thermo Fisher Scientific | 28906 |
| Glycine | Sigma-Aldrich | G7126 |
| cOmplete™ Protease inhibitor cocktail | Sigma-Aldrich | 4693132001 |
| Halt protease and phosphatase inhibitor cocktail | Thermo Fisher Scientific | 78440 |
| Tris-HCl pH 8.0 | Ambion | AM9855G |
| EDTA | Ambion | AM9260G |
| NaCl | Ambion | AM9760G |
| NP-40 | Abcam | ab142227 |
| SDS | Invitrogen | 15553027 |
| DpnII and buffer | New England Biolabs | R0543S |
| Biotin-dATP | Jena Bioscience | NU-835-BIO14-L |
| dCTP/dGTP/dTTP | Invitrogen | 10217016 / 10218014 / 10219012 |
| Klenow | New England Biolabs | M0210S |
| T4 DNA Ligase and buffer | New England Biolabs | M0202S |
| UltraPure™ BSA | Invitrogen | AM2616 |
| Tris-HCl pH 7.5 | Invitrogen | 15567027 |
| Proteinase K | Invitrogen | 25530049 |
| ChIP DNA Clean & Concentrator kit | Zymo Research | D5205 |
| Streptavidin C-1 beads | Invitrogen | 65001 |
| MgCl<sub>2</sub> | Ambion | AM9530G |
| Dimethylformamide | Sigma-Aldrich | 227056 |
| Adapter-loaded Tn5 transposase | In-house preparation from pET28-N-Tn5 EK/LP | N/A |
| NEBNext<sup>®</sup> Ultra<sup>™</sup> II Q5<sup>®</sup> Master Mix | New England Biolabs | M0544S |
| Qubit dsDNA, HS | Invitrogen | Q32851 |
| SPRISelect beads | Beckman Coulter | B23317 |
| RNAClean XP | Beckman Coulter | A63987 |
| Buffer EB | QIAGEN | 19086 |
| RNeasy Plus Mini Kit | QIAGEN | 74134 |
| TruSeq Stranded Total RNA Library Prep Kit | Illumina | 20020596 |
| Ribo-Zero Plus rRNA Depletion | Illumina | 20037135 |
| Sodium deoxycholate | Bioworld | 40430018-2 |
| BCA Protein Assay Kit | Thermo Fisher Scientific | 23227 |
| NuPAGE<sup>™</sup> LDS Sample Buffer | Invitrogen | NP0007 |
| β-mercaptoethanol | Sigma-Aldrich | M6250 |
| NuPAGE<sup>™</sup> 4-12% Bis-Tris Protein Gels | Invitrogen | NP0322BOX |
| NuPAGE<sup>™</sup> MOPS SDS Running Buffer | Invitrogen | NP0001 |
| iBlot<sup>™</sup> 2 Transfer Stacks | Invitrogen | IB23002 |
| PBST | Thermo Fisher Scientific | 28352 |
| 7mm coverslip | Warner Instruments | CS-7R |
| 12mm coverslip | Warner Instruments | CS-12R |
| 96-well imaging plate | Azenta Life Sciences | MGB096-1-2-LG-L |
| Vacuum suction pen | Jiayouy/Amazon | ASIN: B09LLV9CW2 |

**Supplementary Data 1 | Nuclear protein-focused compound library and hiFISH-96 screening results.** This file lists the 43 compounds included in the focused screen with their annotated targets, compound classes, treatment concentrations, and hiFISH-96-derived measurements of ecDNA organisation. The primary screening metric was ecDNA dispersal, quantified as mean foci per cell. Thirty-nine compounds were evaluated for their ability to increase ecDNA dispersal relative to DMSO-treated controls; four compounds were excluded because substantial cell death prevented reliable image acquisition. Additional spatial metrics, including peak location, defined as the radial position of maximum enrichment, and right 75% width, defined as the distance from the peak to the position on the right-hand side of the radial profile at which enrichment decreases to 75% of the peak value, are provided where applicable.

## References

1. Kim, H. et al. Extrachromosomal DNA is associated with oncogene amplification and poor outcome across multiple cancers. Nat Genet 52, 891–897 (2020).

2. Luebeck, J. et al. Extrachromosomal DNA in the cancerous transformation of Barrett’s oesophagus. Nature 616, 798–805 (2023).

3. Lange, J. T. et al. The evolutionary dynamics of extrachromosomal DNA in human cancers. Nat Genet 54, 1527–1533 (2022).

4. Bailey, C. et al. Origins and impact of extrachromosomal DNA. Nature 635, 193–200 (2024).

5. Yan, X., Mischel, P. & Chang, H. Extrachromosomal DNA in cancer. Nat. Rev. Cancer 24, 261–273 (2024).

6. Hung, K. L. et al. ecDNA hubs drive cooperative intermolecular oncogene expression. Nature 600, 731–736 (2021).

7. Turner, K. M. et al. Extrachromosomal oncogene amplification drives tumour evolution and genetic heterogeneity. Nature 543, 122–125 (2017).

8. Wu, S. et al. Circular ecDNA promotes accessible chromatin and high oncogene expression. Nature 575, 699–703 (2019).

9. Hung, K. L. et al. Coordinated inheritance of extrachromosomal DNAs in cancer cells. Nature 635, 201–209 (2024).

10. Nichols, A. et al. Chromosomal tethering and mitotic transcription promote ecDNA nuclear inheritance. Mol. Cell 85, 2839–2853.e8 (2025).

11. Li, Z. et al. Development and Characterization of a Wee1 Kinase Degrader. Cell Chem. Biol. 27, 57–65.e9 (2020).

12. Hirai, H. et al. Small-molecule inhibition of Wee1 kinase by MK-1775 selectively sensitizes p53-deficient tumor cells to DNA-damaging agents. Mol. Cancer Ther. 8, 2992–3000 (2009).

13. Rock, K. L. et al. Inhibitors of the proteasome block the degradation of most cell proteins and the generation of peptides presented on MHC class I molecules. Cell 78, 761–771 (1994).

14. Rorà, A. G. L. di, Cerchione, C., Martinelli, G. & Simonetti, G. A WEE1 family business: regulation of mitosis, cancer progression, and therapeutic target. J. Hematol. Oncol. 13, 126 (2020).

15. Ando, R., Sakaue-Sawano, A., Shoda, K. & Miyawaki, A. Two coral fluorescent proteins of distinct colors for sharp visualization of cell-cycle progression. Cell Struct. Funct. 48, 135–144 (2023).

16. Thangaretnam, K. et al. WEE1 inhibition in cancer therapy: Mechanisms, synergies, preclinical insights, and clinical trials. Crit. Rev. Oncol. Hematol. 211, 104710 (2025).

17. Sankar, V. et al. Genetic elements promote retention of extrachromosomal DNA in cancer cells. Nature 649, 152–160 (2026).

18. Beck, H. et al. Cyclin-Dependent Kinase Suppression by WEE1 Kinase Protects the Genome through Control of Replication Initiation and Nucleotide Consumption. Mol. Cell. Biol. 32, 4226–4236 (2012).

19. Vassilev, L. T. et al. Selective small-molecule inhibitor reveals critical mitotic functions of human CDK1. Proc. Natl. Acad. Sci. 103, 10660–10665 (2006).

20. Hendzel, M. J. et al. Mitosis-specific phosphorylation of histone H3 initiates primarily within pericentromeric heterochromatin during G2 and spreads in an ordered fashion coincident with mitotic chromosome condensation. Chromosoma 106, 348–360 (1997).

21. House, N. C. et al. Profiling the Activity of the Potent and Highly Selective CDK2 Inhibitor BLU-222 Reveals Determinants of Response in CCNE1-Aberrant Ovarian and Endometrial Tumors. Cancer Res. 85, 1297–1309 (2025).

22. Pardee, A. B. A Restriction Point for Control of Normal Animal Cell Proliferation. Proc. Natl. Acad. Sci. 71, 1286–1290 (1974).

23. Pardee, A. B. G1 Events and Regulation of Cell Proliferation. Science 246, 603–608 (1989).

24. Cremer, T. & Cremer, M. Chromosome Territories. Cold Spring Harb. Perspect. Biol. 2, a003889 (2010).

25. Guin, K., Keikhosravi, A., Chari, R., Pegoraro, G. & Misteli, T. Orderly mitosis shapes interphase genome architecture. 10.7554/elife.108410 (2026).

26. Wang, R., Yang, J. F., Ho, F., Robertson, E. S. & You, J. Bromodomain-Containing Protein BRD4 Is Hyperphosphorylated in Mitosis. Cancers 12, 1637 (2020).

27. Hernandez-Verdun, D. Assembly and disassembly of the nucleolus during the cell cycle. Nucleus 2, 189–194 (2011).

28. Faber, G. P., Nadav-Eliyahu, S. & Shav-Tal, Y. Nuclear speckles – a driving force in gene expression. J. Cell Sci. 135, jcs259594 (2022).

29. Lång, A., Lång, E. & Bøe, S. O. PML Bodies in Mitosis. Cells 8, 893 (2019).

30. Rai, A. K., Chen, J.-X., Selbach, M. & Pelkmans, L. Kinase-controlled phase transition of membraneless organelles in mitosis. Nature 559, 211–216 (2018).

31. Li, Y.-D. et al. Template-assisted covalent modification underlies activity of covalent molecular glues. Nat. Chem. Biol. 20, 1640–1649 (2024).

32. Hassan, M. M. et al. Exploration of the tunability of BRD4 degradation by DCAF16 trans-labelling covalent glues. Eur. J. Med. Chem. 279, 116904 (2024).

33. Nathanson, D. A. et al. Targeted Therapy Resistance Mediated by Dynamic Regulation of Extrachromosomal Mutant EGFR DNA. Science 343, 72–76 (2014).

34. Finn, E., Misteli, T. & Pegoraro, G. High-throughput DNA FISH (hiFISH). In Spatial Genome Organization, Methods and Protocols. (ed. Sexton, T.) Methods Mol. Biol. 2532, 245– 274 (2022).

35. Stringer, C., Wang, T., Michaelos, M. & Pachitariu, M. Cellpose: a generalist algorithm for cellular segmentation. Nat. Methods 18, 100–106 (2021).

36. Berg, S. et al. ilastik: interactive machine learning for (bio)image analysis. Nat. Methods 16, 1226–1232 (2019).

37. Bragantini, J. et al. Ultrack: pushing the limits of cell tracking across biological scales. Nat. Methods 22, 2423–2436 (2025).

38. Caillat, C. & Perrakis, A. Cdt1 and geminin in DNA replication initiation. Subcell Biochem 62, 71–87 (2012).

39. Mumbach, M. R. et al. HiChIP: efficient and sensitive analysis of protein-directed genome architecture. Nat. Methods 13, 919–922 (2016).

40. Servant, N. et al. HiC-Pro: an optimized and flexible pipeline for Hi-C data processing. Genome Biol. 16, 259 (2015).

41. Dobin, A. et al. STAR: ultrafast universal RNA-seq aligner. Bioinformatics 29, 15–21 (2012).

42. Frankish, A. et al. GENCODE: reference annotation for the human and mouse genomes in 2023. Nucleic Acids Res. 51, D942–D949 (2023).

43. Liao, Y., Smyth, G. K. & Shi, W. featureCounts: an efficient general purpose program for assigning sequence reads to genomic features. Bioinformatics 30, 923–930 (2014).

44. Love, M. I., Huber, W. & Anders, S. Moderated estimation of fold change and dispersion for RNA-seq data with DESeq2. Genome Biol. 15, 550 (2014).

45. Subramanian, A. et al. Gene set enrichment analysis: A knowledge-based approach for interpreting genome-wide expression profiles. Proc. Natl. Acad. Sci. 102, 15545–15550 (2005).

46. Fang, Z., Liu, X. & Peltz, G. GSEApy: a comprehensive package for performing gene set enrichment analysis in Python. Bioinformatics 39, btac757 (2023).

47. Liberzon, A. et al. The Molecular Signatures Database Hallmark Gene Set Collection. Cell Syst. 1, 417–425 (2015).

